# A spike-ferritin nanoparticle vaccine induces robust innate immune activity and drives polyfunctional SARS-CoV-2-specific T cells

**DOI:** 10.1101/2021.04.28.441763

**Authors:** Joshua M. Carmen, Shikha Shrivastava, Zhongyan Lu, Alexander Anderson, Elaine B. Morrison, Rajeshwer S. Sankhala, Wei-Hung Chen, William C. Chang, Jessica S. Bolton, Gary R. Matyas, Nelson L. Michael, M. Gordon Joyce, Kayvon Modjarrad, Jeffrey R. Currier, Elke Bergmann-Leitner, Allison M.W. Malloy, Mangala Rao

## Abstract

Potent cellular responses to viral infections are pivotal for long -lived protection. Evidence is growing that these responses are critical in SARS -CoV-2 immunity. Assessment of a SARS -CoV-2 spike ferritin nanoparticle (SpFN) immunogen paired with two distinct adjuvants, Alhydrogel® (AH) or Army Liposome Formulation containing QS-21 (ALFQ) demonstrated unique vaccine evoked immune signatures. SpFN+ALFQ enhanced recruitment of highly activated classical and non -classical antigen presenting cells (APCs) to the vaccine-draining lymph nodes of mice. The multifaceted APC response of SpFN+ALFQ vaccinated mice was associated with an increased frequency of polyfunctional spike -specific T cells with a bias towards T_H_1 responses and more robust SARS-CoV-2 spike-specific recall response. In addition, SpFN+ALFQ induced K^b^ spike_(539-546)_-specific memory CD8^+^ T cells with effective cytolytic function and distribution to the lungs. This epitope is also present in SARS-CoV, thus suggesting that generation of cross-reactive T cells may provide protection against other coronavirus strains. Our study reveals that a nanoparticle vaccine, combined with a potent adjuvant, generates effective SARS-CoV-2 specific innate and adaptive immune T cell responses that are key components to inducing long-lived immunity.

**One Sentence Summary:** SpFN vaccine generates multifactorial cellular immune responses.

## INTRODUCTION

Coronaviruses (CoV) are positive-sense, single stranded RNA viruses that cause varying disease pathologies ranging from common cold symptoms to acute respiratory distress syndrome, as well as, gastrointestinal symptoms *(1-3)*. SARS-CoV-2, the causative agent of COVID-19, represents the seventh CoV to be isolated from humans, and is the third to cause severe disease *(4)*. The rapid and unparalleled spread of SARS -CoV-2 into a global pandemic has driven an urgent need for rapidly deployable and scalable vaccine platforms. A myriad of vaccine platforms and approaches have been adopted and developed by government, industry, academic, and non-governmental organizations. Two messenger RNA -based vaccines and an adenovirus vector-based vaccine have been approved for emergency use in the United States *(5)* (https://www.raps.org/news-and-articles/news-articles/2020/3/covid-19-vaccine-tracker). While messenger RNA -based vaccines (*6, 7*) and recombinant adenovirus vectored vaccines (*8-10*) have demonstrated potent efficacy and unprecedented deployment speed, there exists a need for precision vaccine design that may offer improved efficacy to different demographic groups, induce durable immune responses, and provide a multivalent strategy to protect against multiple CoV species and the recent emergence of rapidly evolving immunological variants (*11-13*).

We have recently developed a SARS-CoV-2 sub-unit vaccine based on the ferritin nanoparticle platform (*14*) that displays a pre-fusion stabilized spike on its surface (*15*) and has the potential to fill these current gaps. The stabilized prefusion-spike protein of the Wuhan-Hu-1 strain of SARS-CoV-2 was genetically linked to form a ferritin-fusion recombinant protein, which naturally forms a Spike -Ferritin nanoparticle (SpFN). Ferritin is a naturally occurring, ubiquitous, iron -carrying protein that self -oligomerizes into a 24 -unit spherical particle and is currently being evaluated as a vaccine platform for influenza in two phase 1 clinical trials (NCT03186781, NCT03814720) with two further trials in the recruitment phase for Epstein Barr virus (NCT04645147) and Influenza H10 (NCT04579250).

The spike protein was modified in the following ways to generate a stable spike trimer formation on the ferritin molecule. Two proline residues (K986P, V987P) were introduced to the spike ectodomain, the furin cleavage site was mutated (RRAS to GSAS) (*16*), and the coil-coil interactions were stabilized by mutating the heptad repeat between hinge 1 and 2 (residues 1140-1161). In addition, a short linker to the ferritin molecule was used to exploit the natural three-fold axis, for display of eight spikes. SpFN was then formulated with either o f the two distinct adjuvants to evaluate innate and adaptive T cell responses induced by each of the adjuvants. Our prior experience with Army Liposome Formulation containing the saponin, QS -21(ALFQ) made it a top candidate to compare with one of the most commonly used adjuvants, Aluminum Hydroxide gel (Alhydrogel ^®^) (AH). ALFQ contains two immunostimulants, synthetic MPLA (3D -PHAD^®^) and QS-21 (*17*), the same two immunostimulants that are also present in AS01B, the adjuvant in the highly efficacious licensed herpes -zoster vaccine, Shingrix™ (*18*). However, the liposomes in ALFQ fundamentally differ from those in AS01B, bo th in the phospholipid type and cholesterol content in addition to differences in the amount of 3D -PHAD^®^ and QS-21. Previously, it has been shown in several different models that ALFQ generates well -balanced T_H_1/T_H_2 immunity and protective efficacy (*19-21*).

In this study, we show for the first time the effect of adjuvant design on antigen presenting cell (APC) recruitment to the lymph nodes draining the vaccination site, and its impact on a SARS -CoV-2 vaccine platform. We demonstrate that SpFN formulated with ALFQ (SpFN+ALFQ) compared to SpFN formulated with AH (SpFN+AH) significantly increased recruitment and activation of classical and non -classical APCs. Intriguingly, this was associated with a potent T_H_1-biased cellular response and highly functional spike -specific memory T cells. The APC response to SpFN+ALFQ was characterized by conventional type 1 and type 2 dendritic cells (cDC1 and cDC2) with upregulated costimulatory molecules, in contrast to SpFN+AH. The effective APC response induced by SpFN+ALFQ correlated with differentiation of spike -specific CD4^+^ T cells expressing markers of T_H_1 phenotype. Strikingly, vaccination with SpFN+ALFQ resulted in spike -specific CD8^+^ T cells that established a memory pool. We identified eleven SARS -CoV-2 T cell epitopes in C57BL/6 mice vaccinated with SpFN+ALFQ that mapped to the spike protein. Several of these epitopes are corroborated by recently described findings using an overlapping SARS-CoV-2 peptide library (*22*) following SARS-CoV-2 infection in H2 ^b^ restricted mice. Using an MHC class I tetramer, we identified murine K^b^ restricted SARS-CoV-2 specific memory CD8^+^ T cells recognizing an eight amino acid sequence (amino acids 539 -546; VNFNFNGL) of the SARS-CoV-2 spike protein that is conserved in the SARS -CoV spike protein. The K^b^ spike(539-546)-specific memory CD8^+^ T cells generated by vaccination with SpFN+ALFQ exhibited significantly more cytotoxic activity compared to those identified in the SpFN+AH vaccinated mice. Together these finding demonstrate a novel vaccine platform for SAR-CoV-2 that leverages the innate immune response to induce potent memory-specific antiviral T cells.

## RESULTS

### Spike ferritin nanoparticle (SpFN) adjuvanted with ALFQ (SpFN+ALFQ) induced a more robust and sustained APC response compared to formulation with aluminum hydroxide (SpFN+AH)

Adjuvants are immunostimulants that activate the innate immune system and facilitate vaccine antigen presentation (*23, 24*). To define the ability of distinct adjuvants to engage the innate immune response, we compared our novel vaccine formulations, SpFN+ALFQ with SpFN+AH. Mice were vaccinated at the specified time points and innate immune cells were analyzed from the inguinal and popliteal lymph nodes that drain the vaccinated muscle (dLNs) at days 3 and 5 post vaccination (Fig. 1A). A multiparameter spectral flow cytometry panel (table S1) was used to discriminate APC subsets. T -Distributed Stochastic Neighbor Embedding (tSNE) was used to visualize the high-dimensional datasets (Fig. 1B). The flow cytometric gating strategy is shown in fig. S1, A. Significant differences in the composition of APCs present in the dLNs were seen in mice vaccinated with SpFN+ALFQ compared to SpFN+AH (Fig. 1C, and fig. S1, B and C). The overall number of APCs present in the dLNs was increased 3-7-fold in SpFN+ALFQ vaccinated mice compared to SpFN+AH vaccinated mice, as shown by the pie graphs in Fig. 1C. Interestingly, the number and diversity of APCs began to contract 5 days post vaccination in the SpFN+AH vaccinated mice to levels exhibited in naïve mice. In contrast, mice vaccinated with SpFN+ALFQ exhibited continued expansion of APCs in the dLNs at this time point, demonstrating a more sustained response to the adjuvant formulation, ALFQ (Fig. 1C).

**Fig. 1.**
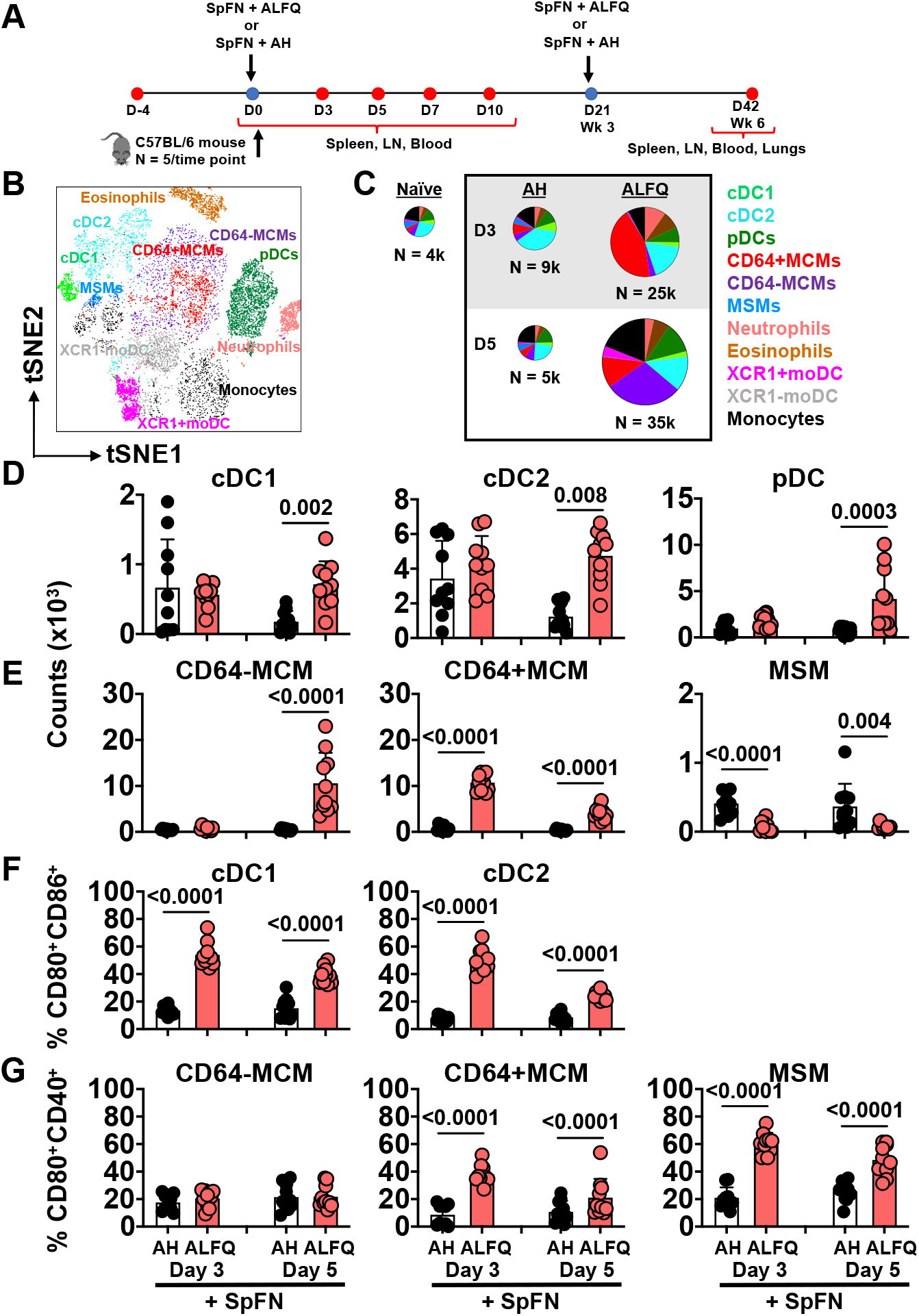
APC recruitment in response to a SARS-CoV-2 vaccine (SpFN) adjuvanted with ALFQ (SpFN+ALFQ) versus AH (SpFN+AH). **(A)** Schematic of vaccination schedule and sample collection. C57BL/6 mice received a prime -boost vaccination regimen with SpFN formulated with either AH (SpFN+AH) or ALFQ (SpFN+ALFQ). At days 0, 3, 5, 7, and 10, spleen, lymph nodes draining the left quadriceps (dLN), and blood were collected. Three weeks (day 42) after the second vaccination (day 21) spleen, mediastinal LN, lungs, and blood were collected. **(B)** The tSNE display of the APC subsets identified by the gating strategy is shown in **Fig. S1A. (C)** Pie representation comparing the number of recruited APCs measured in the dLNs at days 3 and 5 in response to the two vaccine formulations (n = 5/group/time point) or from naÏve mice (n = 4). Eleven APC subsets were identified. Each slice indicates an APC subset; N indicates the average of the total number of APCs recruited in the dLNs. (**D-E)** The number of **(D)** conventional (cDC1, cDC2) and plasmacytoid DCs (pDCs) and **(E)** macrophage subsets in both vaccine groups at both time points. (**F)** Activation of cDCs indicated by CD80 and CD86 and **(G)** macrophage subsets by CD80 and CD40 in both vaccine groups at both time points. Bars indicate mean ± SD. MCM= medullary cord macrophage, MSM=Medullary sinus macrophage, moDC=monocyte derived DC. Experiments were repeated twice and differences between the two groups were analyzed by using non -parametric Mann–Whitney U-test with p ≤ 0.05 considered as statistically significant.

Furthermore, not only were the numbers of APCs increased after SpFN+ALFQ vaccination at days 3 and 5, but classical APCs, such as migratory DCs and lymph node resident APCs, predominated (Fig. 1D). Migratory DCs, which include cDC1 and cDC2s, migrate from tissue sites to dLNs to present antigen and support T cell activation and differentiation and were increased in number in the SpFN+ALFQ vaccinated mice (Fig. 1D). In addition, plasmacytoid DCs (pDCs), that produce and provide type I interferon for DC and T cell activation, were also increased in the dLNs after the priming vaccination with SpFN+ALFQ compared to SpFN+AH (Fig. 1D).

Lymph node resident macrophages play a role in acquisition of antigens through pinocytosis and phagocytosis, and participate in immune regulation. Medullary sinus macrophages (MSM) defined as CD169^+^CD11b^+^F4/80^+^MHCII^int^ CD11c^lo^ and medullary cord macrophages (MCM) defined as CD169 ^-^ CD11b^+^F4/80^+^MHCII^int^ CD11c^lo^CD64^+/-^ (*25, 26*) were the most significantly divergent (p <0.0001) between the two vaccinated groups; higher numbers of MCM were measured in the SpFN+ALFQ vaccinated mice, whereas MSM numbers were significantly higher in the SpFN+AH vaccinated mice, (Fig. 1E). CD64^-^ and CD64^+^MCM were infrequent in numbers on both days 3 and 5 in the SpFN+AH vaccinated mice, while CD64^+^ MCM were present in higher numbers on both days 3 and 5 post-vaccination with SpFN+ALFQ. Interestingly, in the SpFN+ALFQ vaccinated mice, CD64 ^-^MCM, which were infrequent in number on day 3 post-vaccination increased by ∼10,000-fold on day 5 post-vaccination.

Commensurate with increased numbers of classical APCs, the functional activation of these cells was also enhanced by SpFN+ALFQ compared to SpFN+AH. In addition to antigen presentation, costimulatory molecules, such as CD80 and CD86, are expressed by activated DCs and engage T cells, inducing activation rather than anergy in response to ligation with their peptide -MHC complex (*27, 28*). The percentage of cDC1 and cDC2 expressing CD80 and CD86 three days post vaccination was 3 times higher in the SpFN+ALFQ group compared to SpFN+AH (Fig. 1F and fig. S1B). We also measured the activation of macrophage subsets. While the expression of CD80 and CD40 were similarly low in CD64^-^MCMs, the CD64^+^MCMs from SpFN+ALFQ vaccinated mice significantly upregulated CD80 and CD40 by 3 -fold at day 3, and 2-fold at day 5 post priming vaccination compared to the SpFN+AH group (Fig. 1G). In addition, although the MSM numbers were higher with AH as the adjuvant, the expression of costimulatory molecules, CD80 and CD40, was increased in the SpFN+ALFQ vaccinated mice suggesting a higher activation status (Fig. 1G).

Neutrophils, eosinophils, and monocytes have been shown to function as APCs or support antigen-specific T and B cell differentiation in humans and mice (*29-31*). Increased numbers of eosinophils, neutrophils, and monocytes were observed in the dLNs of SpFN+ALFQ vaccinated mice at days 3 and 5 post vaccination (figs. S1, D and E). In addition, a notable proportion of CD11b^+^MHCII^+^ monocyte-like cells were CD11c intermediate, indicative of monocyte-derived DCs (moDCs), and were further differentiated by expression of the chemokine receptor XCR1 (Fig. 1B; fig. 1SA). The XCR1+moDCs were distinct from cDC1, as XCR1+moDCs were MHC -II low, CD11c low, and CD103 negative, and displayed two separate populations by t -SNE. Few XCR1+moDCs were detected in the dLNs of either vaccine group on day 3 post priming vaccination. However, by day 5 post vaccination this moDC subset had significantly increased in the SpFN+ALFQ group suggesting that these cells were involved in acquisition and presentation of the antigen (fig. S1E).

### Effective APC induction by vaccination with SpFN+ALFQ was associated with increased SARS-CoV-2 spike-specific T cells in the dLNs

Next, we analyzed the recruitment of total and spike -specific T cells to the dLNs following vaccination with SpFN+ALFQ or SpFN+AH. T cells from the dLNs were analyzed at days 7 and 10 post priming vaccination for phenotype and cytokine expression, respectively. At day 7 post priming vaccination, the total T cells, as well as the CD4^+^ and CD8^+^ T cell subsets, in the dLNs of SpFN+ALFQ vaccinated mice were significantly higher than those in the SpFN+AH vaccinated mice (Fig. 2A). T cell memory differentiation potential was characterized by the expression of CD62L and CD44 (fig. S2A). In addition to the numeric differences of the T cell subsets, SpFN+ALFQ vaccination resulted in proportionally more effector memory -like (CD62L^-^CD44^+^) CD4^+^ and CD8^+^ T cells. In contrast, SpFN+AH vaccination induced more naïve T cells (CD62L^+^CD44^-^) to the dLNs, although this subset was seen after vaccination in both groups (fig. S2B).

**Fig. 2.**
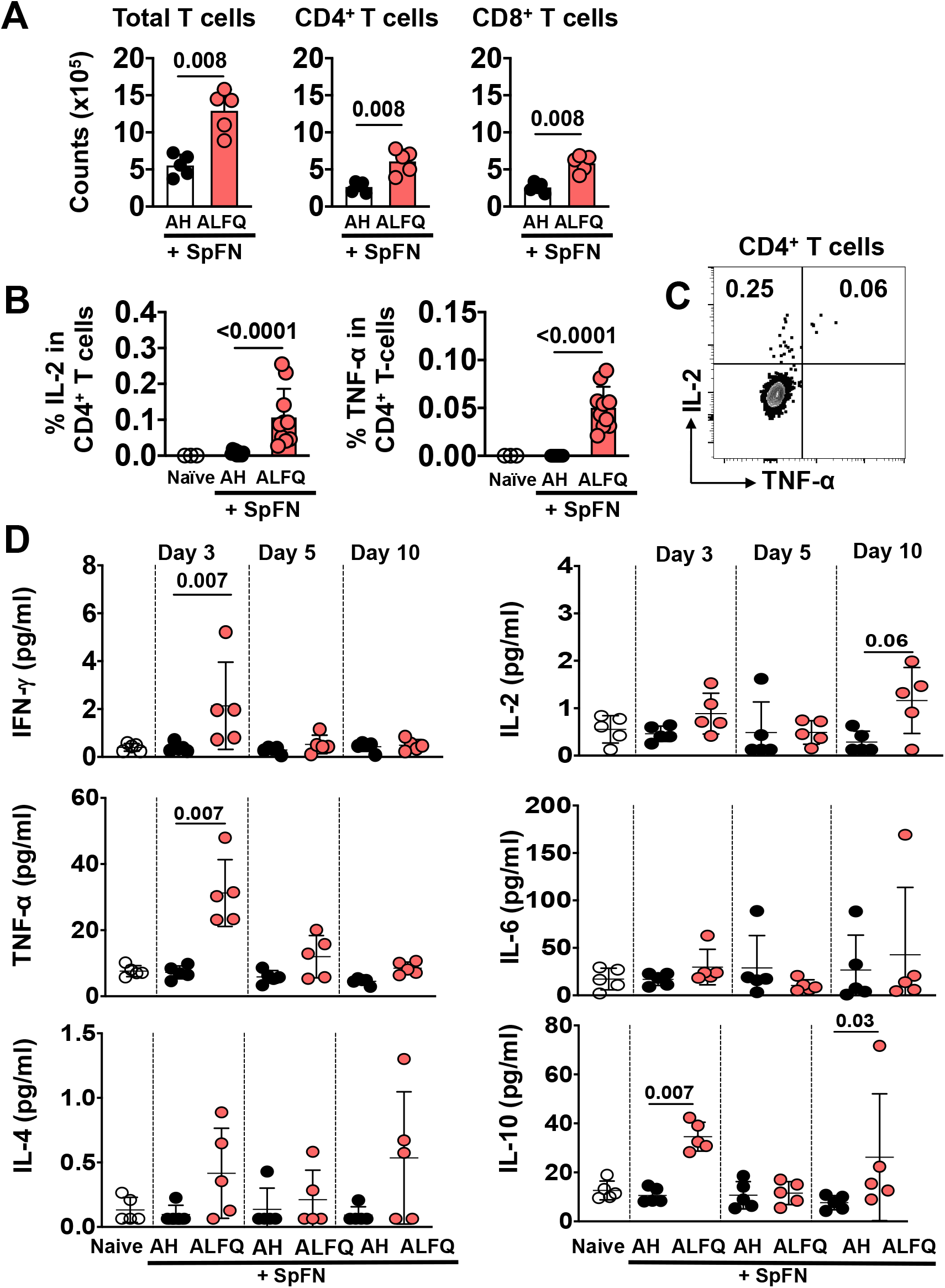
T cell priming in response to SpFN vaccine. **(A)** Magnitude of the T cell response in the dLNs at day 7 post vaccination (n = 5 mice/group/time point). **(B)** Percentage of IL-2 expressing (left panel) and TNF-α expressing (right panel) SARS-CoV-2 spike-specific CD4 ^+^ T cells in the dLNs 10 days post vaccination, following stimulation with spike S1 peptide pool. Bars indicate mean ± SD. **(C)** IL-2 and TNF-α are co-expressed in the SARS-CoV-2 spike-specific CD4 ^+^ T cells, indicating mainly a T _H_ 1 type of response. **(D)** Cytokine profile in the sera of vaccinated mice as determined by multiplex ELISA using Meso Scale Discovery (MSD) platform on days 3, 5, and 10 post -vaccination. Dots represent data from individual mice. Experiments were repeated twice and differences between the two groups were analyzed by using non-parametric Mann–Whitney U-test with p ≤ 0.05 considered as statistically significant.

At day 10 post priming vaccination, SARS -CoV-2 spike-specific T cells were identified by intracellular cytokine staining (ICS) following stimulation with a spike peptide pool (S1) containing the epitopes in the receptor binding domain (RBD). Spike-specific CD4^+^ T cells were readily detected by expression of IL-2 and TNF-αupon peptide stimulation in the dLNs of mice vaccinated with SpFN+ALFQ, and to a much lower extent (p<0.0001) in those of SpFN+AH vaccinated mice (Fig. 2B). IL-2 and TNF-α were co-expressed in a subset of the CD4^+^ T cells (Fig 2C), which were primarily CD44^+^CD62L^-^CCR7^-^ effector memory CD4^+^ T cells (fig. S2C). The intracellular cytokine responses measured from the spike -specific T cells indicate that the CD4^+^ T cell responses were predominantly directed toward T_H_1 (TNF-α and IL-2) rather than a T_H_2 profile. To further confirm our findings, multiplex cytokine analysis was performed with the sera of mice after priming vaccination on days 3, 5 and 10. The T_H_1 cytokines (IFN-γ and TNF-α), as well as IL -10 were significantly higher in the peripheral blood of the SpFN+ALFQ vaccinated mice (Fig. 2D) indicating that the cytokine profile was skewed towards a T_H_1 profile, which was consistent with the measurement of intracellular cytokine expression in the SpFN+ALFQ vaccinated mice.

### Early expansion of SARS-CoV-2 spike-specific CD4^+^ and CD8^+^ T cells following priming vaccination with SpFN+ALFQ

Maturation and activation of APCs, especially DCs, in the dLNs is a prerequisite for the induction of antigen-specific T cell responses and maintenance of long -term memory responses. Since SpFN+ALFQ triggered more robust APC activation along with significantly higher percentages of functional antigen -specific CD4^+^ T cells in the dLNs, we next sought to quantify and characterize the kinetics of vaccine -induced T cell expansion. We therefore evaluated the phenotype and functional CD4^+^ and CD8^+^ T cell responses in the spleen at days 5 and 10 post vaccination. Cells were stimulated with SARS -CoV-2 spike-specific peptides from the S1 and S2 peptide pools from JPT, followed by surface and ICS and measurement by flow cytometry. Although there were no significant differences in the frequency of total T cells, CD4^+^ and CD8^+^ T cells at day 5 and day 10 post vaccination (fig. S3A), we did observe significant differences in the SARS -CoV-2 spike-specific cytokine expressing CD4^+^ and CD8^+^ T cells. CD4^+^ T cells expressing IL-2 (p=0.008) and IFN-γ (p=0.03) were significantly higher in the ALFQ group compared to the AH group at day 10 (Fig. 3, A to C). In addition, CD8^+^ T cells secreting IL-2 (Day 5: p=0.01; Day 10: p=0.03, fig. S3B), IFN-γ (Day 5: p=0.007; Day 10: p=0.03, Fig. S3C) and TNF-α (Day 5: p=0.007; Day 10: p=0.01, fig. S3D) were higher in SpFN+ALFQ vaccine group on both days 5 and 10 post-vaccination (Fig. 3, A to C). The frequency of cytokine-positive cells was generally higher in the CD8^+^ T cell population than the CD4^+^ T cell population at day 10 compared to day 5 post-vaccination (Fig. 3, A to C).

**Fig. 3.**
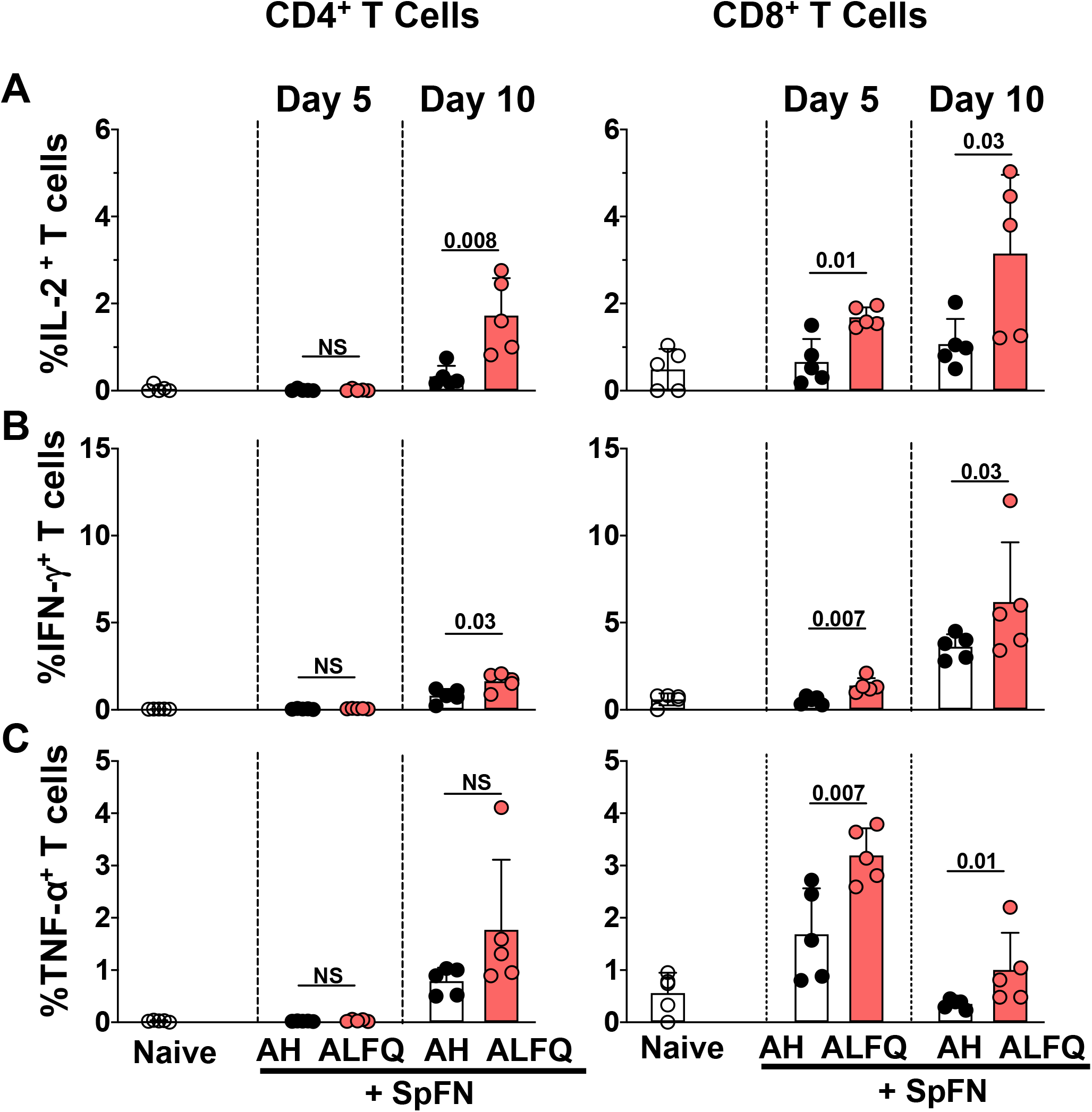
Splenic T cell responses in mice vaccinated with SpFN vaccine. **(A-C)** Flow cytometry-based characterization of mouse splenocytes at days 5 and 10 post-vaccination measuring SARS-CoV-2 spike-specific intracellular cytokine responses in CD4 ^+^ and CD8^+^ T cells **(A)** IL-2-, (**B**) IFN-γ- and (**C**) TNF-α-secreting T cells. Dots represent data from individual vaccinated or na Ïve mice (n= 5/group). Bars represent mean ± SD. (error bar). Statistical significance between the groups was determined using non-parametric Mann–Whitney U-test with p ≤ 0.05 considered as statistically significant. The gating strategy applied for the evaluation of flow-cytometry-acquired data presented in this figure is provided in **Fig. S4**.

### SARS-CoV-2 spike-specific T cells localize with higher frequency and a T_H_1-biased cytokine profile to the lung draining lymph nodes of SpFN+ALFQ vaccinated mice following a prime-boost vaccination

To determine whether an intramuscular vaccination could prime SARS -CoV-2-specific memory T cells to home to tissue sites relevant to potential infection, the T cell response in the mediastinal lymph nodes were measured three weeks following a booster vaccination. The gating strategy for the identification of SARS -CoV-2 spike-specific CD4^+^ and CD8^+^ T cells is shown in fig. S2D. Spike-specific CD4^+^ and CD8^+^ T cells expressed IFN-γ, TNF-α, IL-17A (Fig. 4A) and IL-10 (Fig. 4B) in the SpFN+ALFQ vaccine group. The IFN-γ expressing CD4^+^ and CD8^+^ T cells exhibited activated effector memory markers (CD69^+^ CD44^+^ and CD62L^-^ CD103^-^; fig. S2, E and F). Furthermore, IFN-γ and TNF-α were dominant and co-expressed by both spike-specific CD4^+^ and CD8^+^ T cell in the SpFN+ALFQ vaccine group (Fig. 4C).

**Fig. 4.**
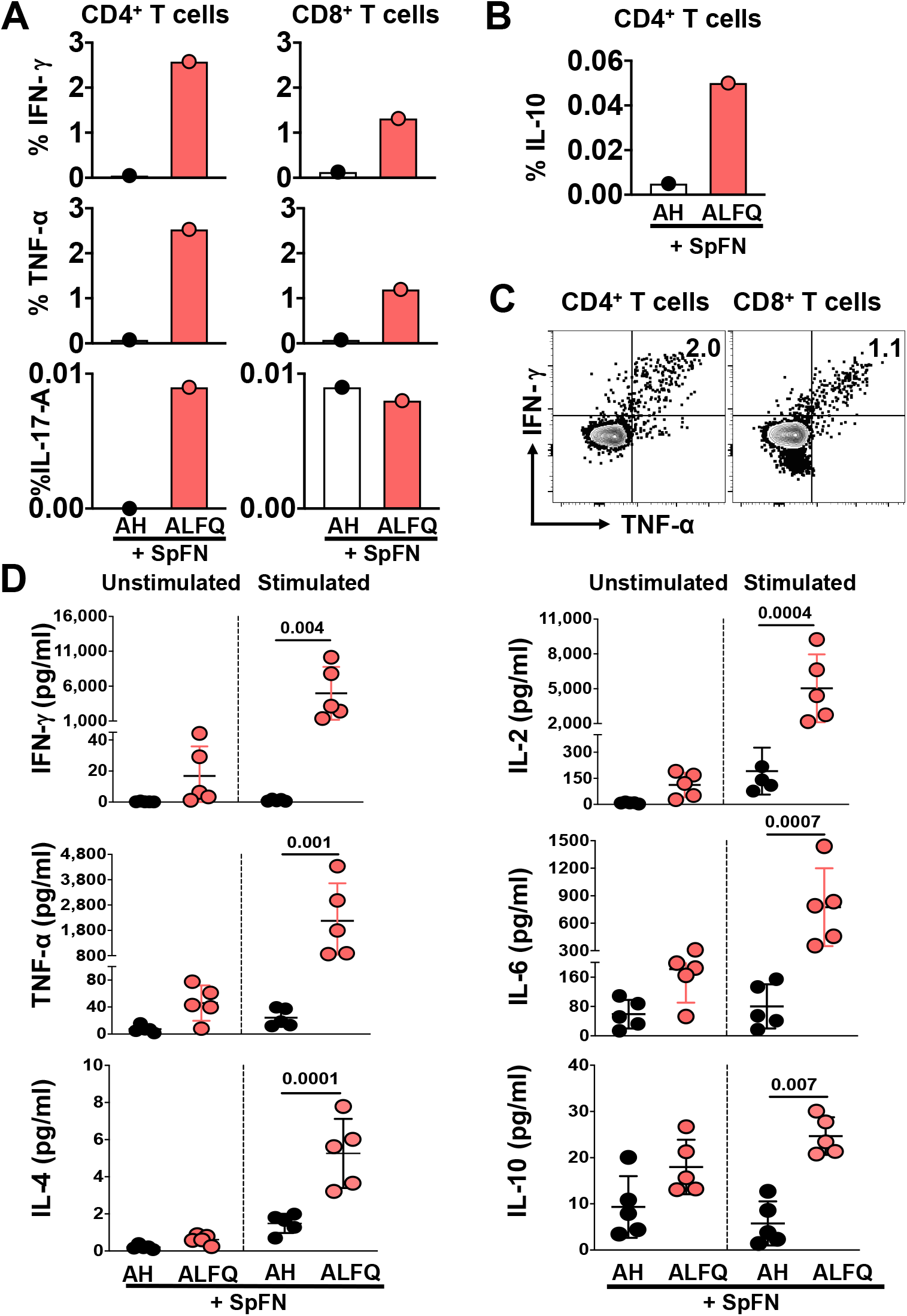
Cytokine responses in the mediastinal lymph nodes and spleens of mice after SpFN prime -boost vaccination. **(A-B)** Percentage of SARS-CoV-2 spike-specific T cells in the mediastinal lymph nodes three weeks post prime -boost vaccination **(A)** Percentage of CD4^+^ and CD8^+^ T cells expressing IFN-γ, TNF-α, IL-17A and **(B)**, CD4^+^ T cells expressing IL -10. Each sample is representative of the pooled lymph nodes within the vaccine group. Bars indicate mean ± SD. **(C)** SARS-CoV-2 spike-specific CD4 ^+^ and CD8^+^ T cells in the mediastinal lymph nodes following prime -boost vaccination with SpFN+ALFQ co-express IFN-γ and TNF-α upon peptide stimulation. **(D)** Splenocytes were stimulated with SARS -CoV-2 spike-specific peptides and cytokines in the culture supernatants were measured by multiplex ELISA using MSD platform. Dots represent data from i ndividual mice (n=5/group). Differences between the two groups were analyzed by using non -parametric Mann–Whitney U-test with p ≤ 0.05 considered as statistically significant.

The profile of pro-inflammatory cytokines was also compared in the spleen at 3 weeks following booster vaccination to track the differentiation of antiviral T cell responses. Splenocytes were stimulated with pooled SARS-CoV-2 spike peptides spanning the S1 and S2 subunits and the cytokine profiles were assessed. Assessing the two vaccination groups, IFN-γ, TNF-α, IL-2 and IL-6 levels were significantly increased in SpFN+ALFQ vaccinated mice compared to mice that received SpFN+AH vaccination (Fig. 4D). The IL-4 and IL-10 levels were also elevated in the SpFN+ALFQ vaccinated group, although the magnitude of increase was much lower in comparison to the T_H_1 cytokines (Fig. 4D).

### Prime-boost vaccination with SpFN+ALFQ resulted in a more robust SARS -CoV-2 spike-specific T cell response

To assess the durability of T cell responses, we further characterized the T cell phenotypes and functional responses in the spleen at three weeks post booster vaccination. Upon evaluating the total T cell response, we observed an expansion of CD8^+^ T cells compared to CD4^+^ T cells as measured by a decrease in the total CD4^+^ (p=0.002) and an increase in the total CD8^+^ T cells (p=0.0006) in the SpFN+ALFQ compared to SpFN+AH vaccination group (Fig. 5, A and B). To assess the frequency of cytokine secreting CD4^+^ and CD8^+^ T cells, splenocytes were stimulated *ex-vivo* with SARS-CoV-2 spike peptides spanning the S1 or S2 subunits. The flow gating strategy used to determine the frequency of cytokine secreting CD4^+^ and CD8^+^ T cells is represented in **f**ig. S4. Our data revealed that the vaccine boost dramatically increased the frequency of S ARS-CoV-2 spike S1-specific IL-2 (CD4^+^: p= 0.004; CD8^+^: p=0.0005, Fig. 5C), IFN-γ (CD4^+^: p= 0.005; CD8^+^: p=0.0005, Fig. 5D) and TNF-α (CD4^+^: p= 0.0007; CD8^+^: p=0.0005, Fig. 5E) secreting CD4^+^ and CD8^+^ T cells, while no differences were observed in frequency of IL-4 secreting cells (data not shown). The booster dose of SpFN+ALFQ resulted in a further expansion of cytokine producing T cells above the priming response at day 5 and/or day 10. This suggested that the two-dose regimen of the SpFN+ALFQ vaccine formulation improved the generation and differentiation of SARS -CoV-2-specific T cell memory responses. Importantly, the S1 peptide pool containing the RBD and NTD portions of the spike protein produced significantly greater cytokine expression compared to the minimal responses generated by the S2 peptide pool (data not shown). Therefore, we focused further functional assays on this portion of spike that contains both conserved peptide residues and peptides unique to SARS-CoV-2.

**Fig. 5.**
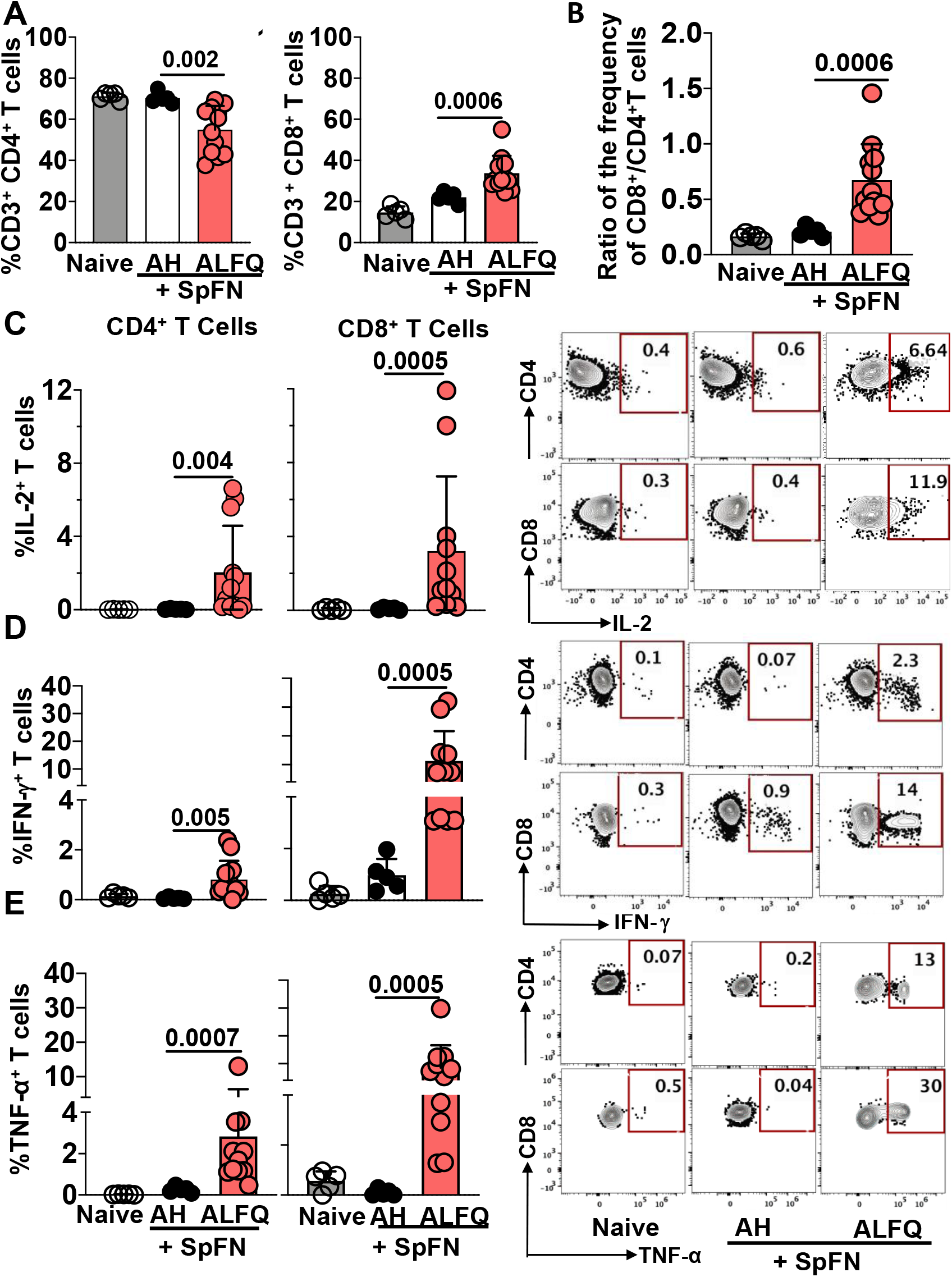
T cell responses in splenocytes of mice in response to SpFN vaccine at week 6 following prime-boost vaccination. **(A)** Frequency of total CD4^+^ and CD8^+^ T cells. **(B)** Ratio of the frequency of total CD8^+^/CD4^+^ T cells. Bar graphs and representative flow plots demonstrating cytokine **(C)** IL-2 **(D)** IFN-γ and **(E)** TNF-α secreting CD4^+^ and CD8^+^ T cells. Dots represent data from individual mice (naive: n =5; AH: n= 5; ALFQ: n= 11, from two independent experiments). Bars represent mean ± SD. (error bar). Statistical significance between the groups was determined using non -parametric Mann–Whitney U-test with p ≤ 0.05 considered as statistically significant.

### T cell responses induced by SpFN+ALFQ are focused to the S -1 domain of the SARS-CoV-2 spike protein

Next, we sought to characterize the peptides of the spike protein targeted by the vaccine induced T cell response. Our data demonstrated that the cytokine response to SARS-CoV-2 was dominated by IFN-γ, therefore an IFN-γ-specific ELISpot assay was used to screen 52 peptide pools composed of 315 peptides as this methodology requires fewer cells and has a higher sensitivity than other measurement assays. Peptides comprised the full length of the spike protein, represented with its major sub -units and domains (S-1 domain containing the NTD, RBD, S-2 domain containing FP, HR1 and HR2) in Fig. 6A. Peptides were pooled in a matrix scheme to allow the high -throughput screening and identification of potential epitopes (Fig. 6B, and fig. S5). Most of the measured reactivity to the peptide pools was focused to the NTD and RBD within S -1 domain of the spike protein with the highest responses in pools 4 and 5, and 19 (Fig. 6C). The results revealed one epitope, VNFNFNGL (aa 539 -546) that was not predicted by the NetMHCPan 4.1 EL prediction tool (iedb.org), but was previously reported (*22*) in mice transfected with human ACE2 mRNA and subsequent infected with SARS -CoV-2, as well as defined in SARS-CoV (*32*). This epitope is conserved between the two viruses. Comparing the results from the NetMHCPan 4.1EL results with the IFN-γ ELISpot results suggest that all epitopes are either H2K^b^ or H2D^b^ restricted and none of them represented CD4^+^ epitopes.

**Fig. 6.**
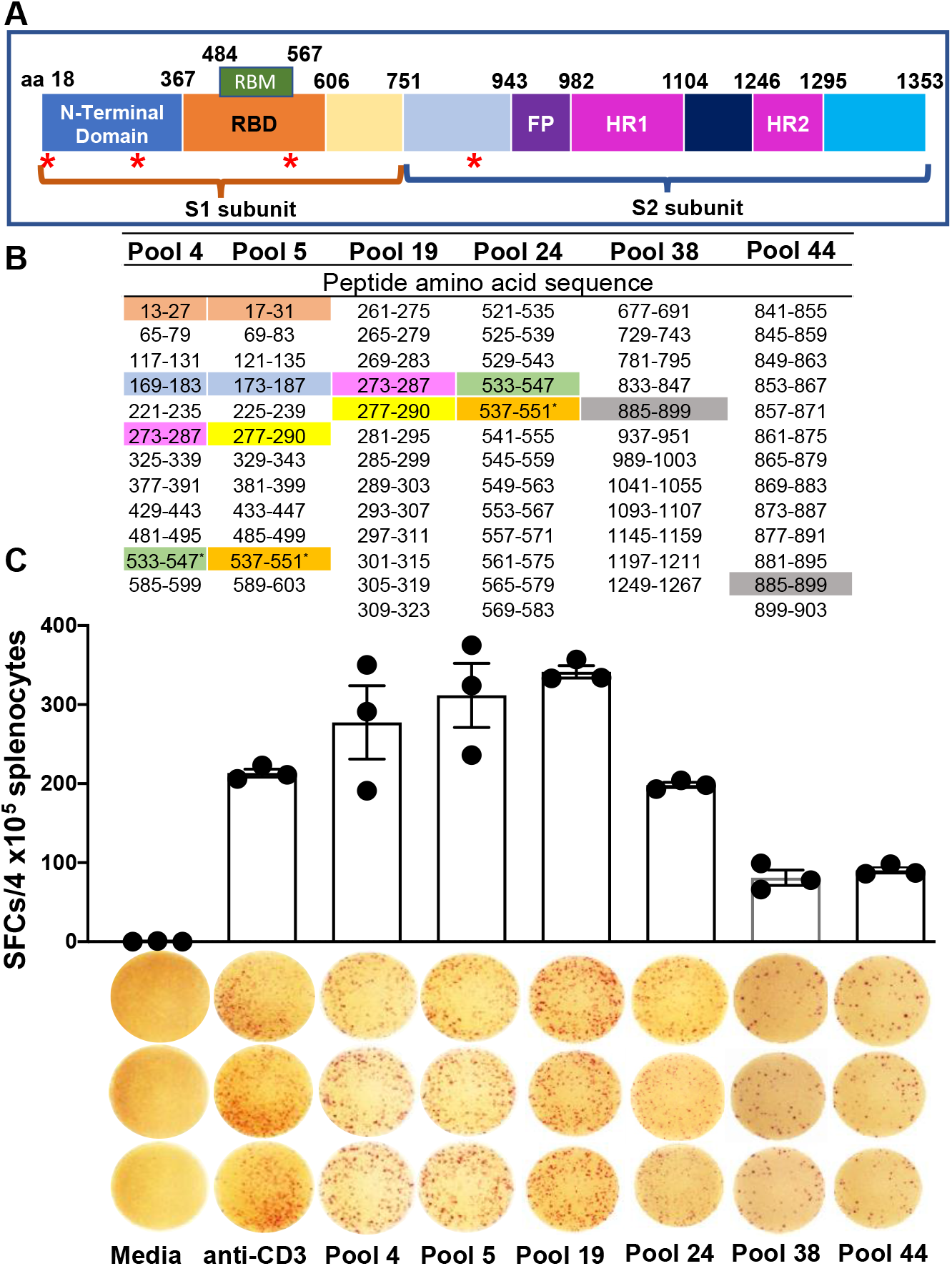
Epitope mapping data in the splenocytes of SpFN vaccinated C57BL/6 mice at week 6 following prime-boost vaccination. **(A)** Schematic representation of the SARS-CoV-2 spike protein showing the different domains with amino acid (aa) numbers. Red asterisk denotes the identified H2K^b^/D^b^-restricted epitopes. **(B)** Amino acid residues of SARS -CoV-2 glycoprotein peptide pools. Matrix pools (4, 5, and 38) and overlapping peptide pools of 15-mer peptides overlapping by 11 aa (19, 24, and 48) containing 315 peptides across the entire glycoprotein were tested in the ELISpot assay. Reactivity to specific matching sequences in pools are highlighted with different matching colors and represent the predicted epitopes for H2K^b^/D^b^ (iedb.org) or predicted epitopes without ELISpot responses (blue highlight; hypothetical epitope). The overlapping peptide pool covering aa 169-1187 (pool 17) did not show any reactivity (data not shown); aa sequences 13-27 (pool 4) and 17-31 (pool 5) have been highlighted in color to indicate confirmed epitopes based on ELISpot reactivity to peptide pool 14, which contains overlapping peptides aa 1 -67 with spot counts of 54, 56, 54 (data not shown). **(C)** Number of spike-specific IFN-γ spot forming units stimulated with 52 peptide pools from the Epitope Mapping Peptide Set of the spike protein is presented as scatter plots (mean ± SD/ 4 x 10^5^ splenocytes). ELISpot images of triplicate wells and the stimulants are shown on the X-axis. T cell responses were considered positive when mean spot count exceeded the mean ± 3 SD of the negative control wells. Splenocytes were stimulated *in vitro* with media (negative control), anti-CD3 (positive control), or SARS CoV-2 spike peptide pools 4, 5, 19, 24, 38, and 44.

### Induction of K^b^-spike_(539-546)_-specific polyfunctional CD8^+^ T cell responses following booster vaccination with SpFN+ALFQ

After establishing the strong effector functions of T cells after the prime-boost vaccination strategy, we then enumerated the frequency of antigen specific CD8^+^ T cells with MHC class I tetramer staining. Epitope mapping data revealed SARS-CoV-2 spike aa 539-546 (VNFNFNGL) as the most immunogenic epitope present in peptide pools 4, 5, and 19 (Fig. 6, B and C). MHC class I, K^b^, tetramers presenting the VNFNFNGL epitope (aa 539 -546) along-with two other H2-K^b^ restricted epitopes, GNYNYLYRL (aa 433 -441) and YNYLYRLF (aa 435 -443) from the spike protein, were used to identify antigen-specific CD8^+^ T cells. A H2 -K^d^ restricted tetramer, containing CYGVSPTKL (aa 365 -373) from the spike protein served as a control (table S5). The gating strategy is provided in fig. S6, A and B. Our data revealed that SpFN+ALFQ generated significantly higher K^b^-spike_(539-546)_-specific CD8^+^ T cells (p=0.01) ranging from 2.47% to 28 % (Fig. 7A) compared to 0.21% to 4.2 % for SpFN+AH (Fig. 7B). Based on the tetramer and epitope mapping data, we performed combined ICS and tetramer staining experiments with the K^b^-spike_(539-546)_-specific tetramer, to demonstrate cytokine secretion by tetramer positive CD8^+^ T cells. Our results demonstrate SpFN+ALFQ vaccinated mice generated higher percentages of K^b^-spike_(539-546)_-specific TNF-α^+^ (Fig. 7C; p =0.007), IFN-γ^+^ CD8^+^ T cells (Fig. 7D; p=0.008), and double positive TNF-α^+^IFN-γ^+^ CD8^+^ T cells (Fig. 7E; p=0.007), compared to SpFN+AH vaccinated mice following the booster vaccination. Cytokine polyfunctionality is of major importance for the successful efficacy of a vaccine (*33*). Several studies have shown a strong correlation between the protection level and the induction of high frequencies of polyfunctional T cells co -expressing IFN-γ, TNF-α, and IL-2 (*33*). In order to determine if SpFN+ALFQ generated antigen-specific polyfunctional CD8^+^ T cells, we assessed their frequency post booster vaccination. Functionality was measured based on stimulation with the S1 peptide pool, as our prior data showed minimal responses to the S2 peptide pool by ICS.

**Fig. 7.**
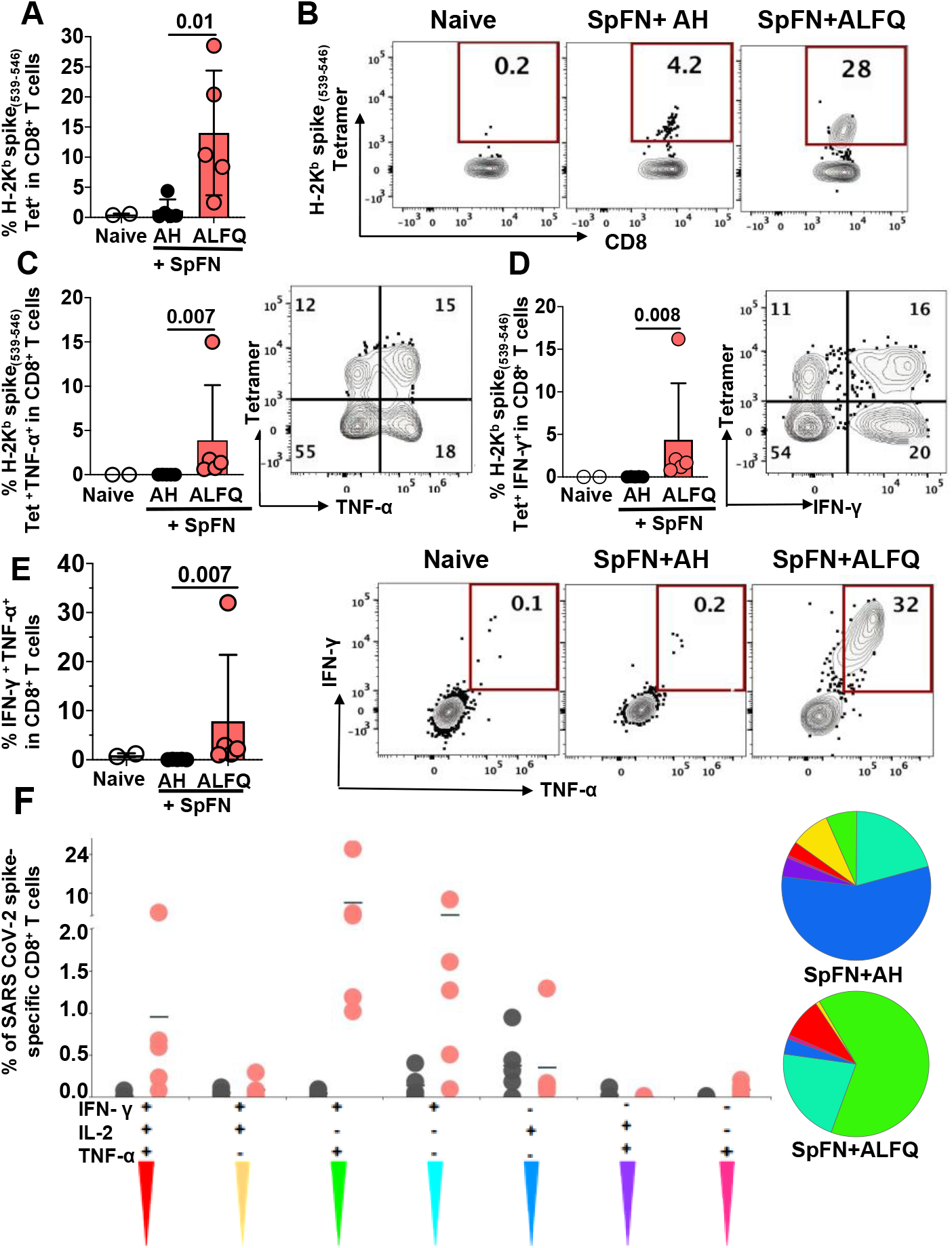
SARS-CoV-2 K^b^ spike _(539-546)_-specific MHC class I restricted CD8^+^ T cells in response to SpFN vaccine following a booster vaccination. **(A)** Frequency of antigen specific CD8^+^ T cells in splenocytes of na Ïve and vaccinated mice as determined by flow cytometry -based SARS-CoV-2 spike _**(539-546)-**_specific tetramer staining. (**B)** Representative flow plots depicting the frequency of K^b^ spike _**(539-546)-**_specific CD8^+^ T cells at week 6 in all the groups. Bar graphs and representative flow plots demonstrating the significantly higher percenta ge of spike _**(539-546)-**_specific-**(C)** TNF-α^+^ and **(D)** IFN-γ^+^ CD8^+^ T cells. **(E)** Bar graph and representative flow plots depicting the significant differences in the frequency of the SARS -Cov-2 spike-specific CD8^+^ T cells concomitantly producing both IFN -γ and TNF-α. Bars represent the mean ± SD. and significant differences between the groups were calculated using non -parametric two-tailed Mann-Whitney U test in Graph pad prism version 8. **(F)** CD8^+^ T cell polyfunctionality: Differences in the ability of CD8^+^ T cells stimulated with SARS-CoV-2 spike peptides to secrete more than one cytokine. Boolean gating was applied to identify all combinations of CD8^+^ T cell effector functions. The pie chart depicts the average proportion of spike -specific CD8^+^ T cells producing all three (IFN -γ, IL-2 or TNF-α), any two or any one cytokine. The graphs show cytokine-secreting peptide-dspecific CD8^+^ T cells for each individual mouse (n = 5 /group) in response to each adjuvant formulation. This analysis was performed using the SPICE software version 5.1(*54*).

SpFN+ALFQ induced a strikingly higher percentage of polyfunctional spike -specific CD8^+^ T cells co-expressing either 3 (IFN-γ, IL-2, and TNF-α) or 2 (IFN-γ, TNF-α) cytokines compared to the SpFN+AH (Fig. 7F). Polyfunctional IFN-γ and TNF-α producing T cells dominated the response from the SpFN+ALFQ vaccinated animals, in contrast to the single cytokine secreting T cell response exhibited by the SpFN+AH group. These data demonstrate the differences in the landscape of T cell responses induced by the two different vaccine formulations.

### K^b^-spike_(539-546)_-specific CD8^+^ T cells from SpFN+ALFQ immunized mice effectively lyse target cells

In order to determine if the induction of a more robust CD8^+^ T cell response in the SpFN+ALFQ vaccinated mice compared to SpFN+AH vaccinated mice translated into improved effector function, we conducted an *in-vitro* killing assay. Target cells were generated by pulsing naïve splenocytes with K^b^-spike_(539-546)_-peptide or left unpulsed and labeled CFSE high or low, respectively. The gating strategy for target CFSE -high and low and tetramer positive effector CD8^+^ T cells is shown in fig. S6C. Cells isolated from the SpFN+ALFQ group demonstrated increased killing of peptide -pulsed target cells compared to SpFN+AH group represented by the ratio of CFSE high:low cells at each dilution (Fig. 8C). Increased CTL activity has been linked with reduced viral titers and inhibition of viral replication (*34*) and is the central function of the CD8^+^ T cells we identified in this study. Representative flow plots further demonstrated an increase in CFSE high population as dilution of effector cells increased (Fig. 8A). K^b^-spike_(539-546)_-specific CD8^+^ T cells present at each dilution were measured. The decrease in cells from the SpFN+ALFQ vaccinated mice corresponded with a decrease in CFSE high cell death (Fig. 8B). Even at a high dilution rate, the cells from SpFN+ALFQ vaccinated animals demonstrated improved cytolytic activity compared to the AH group. Curves for both ALFQ and AH groups were evaluated using Pearson correlation and showed strong positive correlations of 0.823 and 0.888, respectively (Fig. 8C).

**Fig. 8.**
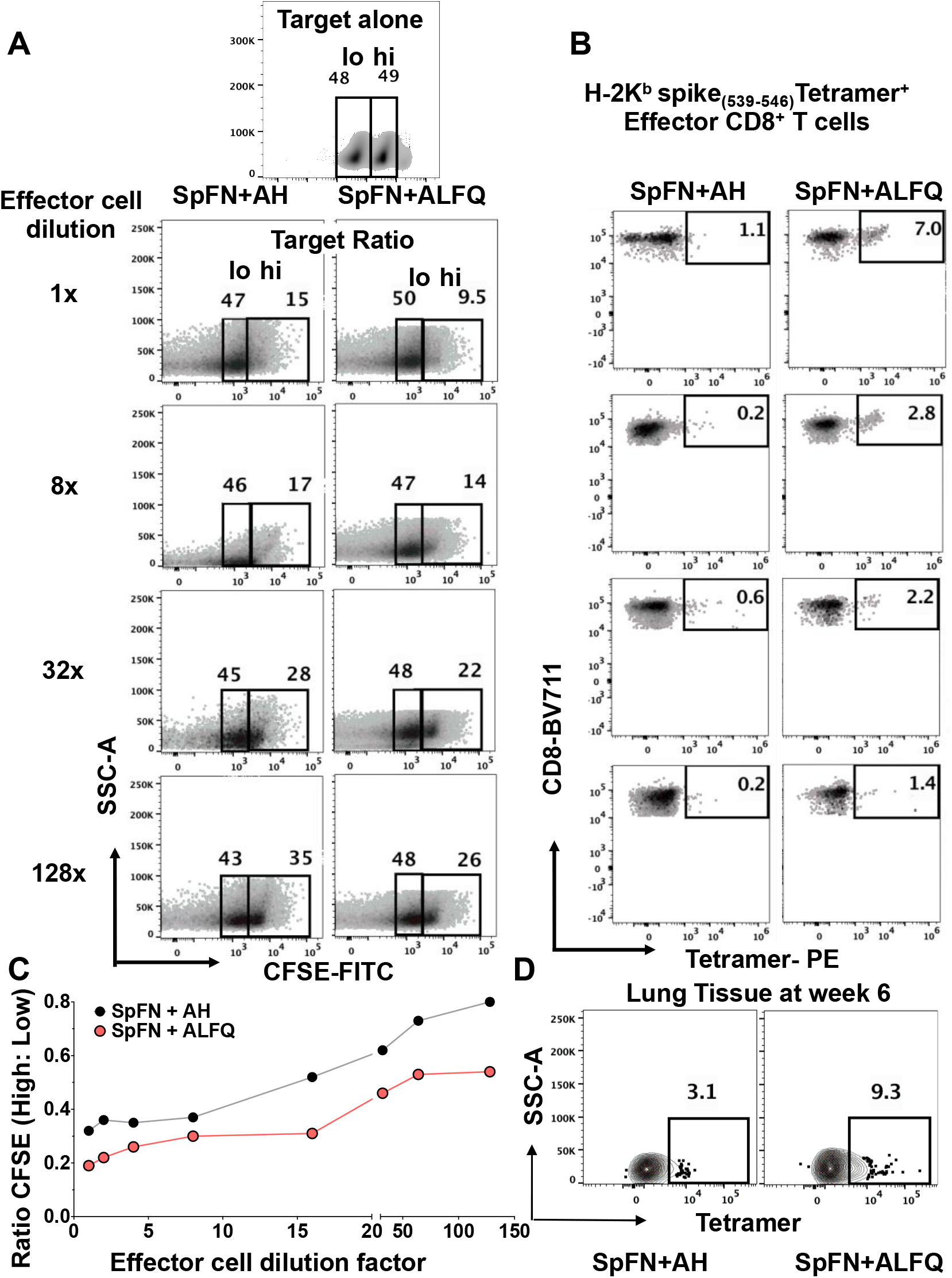
Flow cytometry based CFSE cytotoxic T cell (CTL) assay to measure antigen -specific killing in response to SpFN vaccine at week 6 following prime-boost vaccination. Representative flow cytometry plots depicting the frequency of **(A)** CFSE-high/CFSE-low population and **(B)** effector cells from mice vaccinated with SpFN vaccine adjuvanted with ALFQ versus AH. **(C)** Graph depicting the ratio of CFSE high to low (indicating antigen-specific killing) as a function of effector cell dilution. Each curve represents effector cell dilutions from a single mouse vaccinated with SpFN+ALFQ or SpFN+AH. The curves were evaluated using Pearson correlation coefficient. **(D)** Frequency of SARS-CoV-2 spike _**(539-546)-**_specific tetramer positive CD8^+^ T cells in the lung tissue of SpFN vaccinated mice at week 6 following prime-boost vaccination.

### Prime-boost with SpFN+ALFQ resulted in a large population of spike_(539-546)-_specific memory CD8^+^ T cells in the lungs

Prime-boost with SpFN+ALFQ generated effector memory IFN-γ producing CD4^+^ and CD8^+^ T cells from the mediastinal lymph nodes (Fig. S2, E and F). Therefore, we wanted to determine the distribution of the cytotoxic spike_(539-546)_-specific CD8^+^ T cells at a similar memory time point in the lungs of mice vaccinated with SpFN+ALFQ and SpFN+AH. At 3 weeks following booster vaccination, perfused lungs were processed from both groups of vaccinated mice and stained with K^b^-spike_(539-546)_ tetramer. Tetramer positive CD8^+^ T cells were detected from the lungs of mice in both adjuvant groups (Fig. 8D). Similar to the results from splenocytes, the lungs from the SpFN+ALFQ group exhibited a higher percentage of K^b^-spike_(539-546)-_specific CD8^+^ T cells (9.3%) compared to lungs from the SpFN+AH group (3.1%).

## Discussion

In the present study, we assessed cell -mediated immunity induced by a novel immunogen, SpFN, combined with a potent adjuvant, ALFQ. This candidate vaccine induced robust recruitment of APCs and polyfunctional SARS-CoV-2 spike-specific T cells, and spike epitope-specific cytolytic memory CD8^+^ T cells. The vaccine, SpFN+ALFQ is currently in a phase 1 clinical trial in the United States, sponsored by the U.S. Army (ClinicalTrials.gov Identifier: NCT04784767).

Both humoral and cellular responses have been shown to provide protection against respiratory viral pathogens, such as influenza and RSV (*35, 36*). Moreover, cross-reactive T cells have been shown to protect against a heterologous influenza virus infection in the absence of virus -specific antibodies (*37*). While vaccine evaluation traditionally involves assessment of neutralizing antibodies, currently, the full spectrum of correlates of protection for COVID -19 remain unknown. There are several reports that both CD4^+^ and CD8^+^ T cells specific for SARS-CoV-2 arise during an active infection and during the convalescence period (*38*) and may also provide protection. Resident memory T cells were shown to protect against SARS -CoV in an intranasal mouse vaccine model (*39*), suggesting perhaps a similar important role for T cells in the context of SARS-CoV-2.

Based on the breadth of the observed immune response with SpFN+ALFQ, and protection following challenge with USA-WA1/2020 strain in nonhuman primates (*40*), the induction of broad SARS coronavirus immune responses in mice (unpublished data), a detailed study of the T cell responses induced by SpFN+ALFQ was warranted. The mouse system offers the advantage of defining the *in vivo* mechanisms of innate and adaptive immunity following vaccination with SpFN+ALFQ in comparison with a common and well -established adjuvant AH.

Following vaccination with SpFN+ALFQ, we observed robust and sustained recruitment and activation of classical and nonclassical APCs in the dLNs compared to SpFN+AH. Furthermore, we show that cDC1 and cDC2, in particular, exhibited enhanced upregulation of costimulatory molecules necessary for T cell engagement and differentiation. Interestingly, by thoroughly analyzing the immune response within the dLN of the vaccination site, we discovered that SpFN+ALFQ also recruited more non -classical APCs and monocytes that support recruitment and activation of APCs and T cells that enhance vaccine efficacy (*41*). Moreover, we show that lymph node resident macrophages (*42, 43*) increased in number and functional activation in response to ALFQ+SpFN and may support and optimize plasma cell (*25, 44*) and T cell development (*44*) in response to this vaccine strategy.

Although both vaccine-adjuvant formulations (SpFN+ALFQ and SpFN+AH) recruited naïve T cells, the response elicited by ALFQ was superior and faster at generating or differentiating effector memory phenotype cells than seen with AH. Hence, the adjuvant determines the quality and the quantity of early innate responses which then sets the stage for downstream adaptive immune responses. Furthermore, our results suggest that the effective APC responses induced by SpFN+ALFQ resulted in differentiation of virus -specific CD4^+^ T cells that were biased towards a T_H_1 phenotype (TNF-α and IL-2). Multiplex cytokine data analysis in the sera of SpFN+ALFQ vaccinated mice, post priming, also showed a similar skewing towards a T_H_1 response (IL-2, TNF-α, and IFN-γ). We also noted a significant increase in the secretion of IFN-γ, IL-2, and TNF-α from both CD4^+^ and CD8^+^ T cells from the spleen of SpFN+ALFQ vaccinated mice compared to SpFN+AH in response to spike peptide pools.

Following a booster vaccination, SARS -CoV-2 spike-specific T cell responses were also present in the mediastinal lymph nodes that drain the lungs. A 3 -fold increase in MHC class I restricted K^b^-spike_(539-546)_-specific CD8^+^ T cells was also observed in the lungs of mice vaccinated with SpFN+ALFQ. The importance of T cell surveillance in the lungs cannot be overstated as they are the primary target of SARS-CoV-2 infection. SpFN+ALFQ vaccinated mice exhibited a higher frequency of spike -specific CD4^+^ and CD8^+^ T cells secreting IFN-γ, TNF-α, IL-17A and IL-10, as well as, co-expression of IFN-γ and TNF-α. Multiplex cytokine analysis in the splenocytes conducted at the same time point further validated our ICS results and showed significantly elevated levels of spike -specific T_H_1 cytokines such as IFN-γ, TNF-α, IL-2, and IL-6 compared to vaccination with SpFN+AH. Even though there was an increase in T_H_2 cytokines such as IL-4 and IL-10 with SpFN+ALFQ, the magnitude of the response was minimal in comparison to other T_H_1 cytokines, indicating a strong bias toward secretion of T_H_1 cytokines compared to T_H_2 cytokines. Hence, the two-dose regimen of SpFN+ALFQ vaccine formulation was able to generate strong functional T cell effector and memory responses in the spleen as well as memory T cells in the lungs.

One of the key features of the SpFN+ALFQ cellular response is the early induction of a predominantly T_H_1, and not a T_H_2, response. Vaccine elicited T_H_2 responses have been shown to be detrimental in a SARS-CoV murine infection model (*45-47*). Another important factor is the induction of immune-modulatory cytokines. IFN-γ in particular, is a key cytokine for several antiviral responses (*48, 49*). This cytokine has been shown to act in synergy with type I interferons to inhibit the replication of SARS -CoV (*50*). The robust induction of IFN-γ-producing CD8^+^ T cells in the present study, indicates that SpFN+ALFQ vaccine elicits a favorable cellular immune response and complements the observed strong neutralizing antibody response observed in mice (unpublished) and in nonhuman primates (*40*). The neutralization breadth and the inhibition of ACE -2 binding was maintained with the variants tested (SARS-1, B.1.1.7 {UK} and B.1.351 {SA}) with immune nonhuman primate sera, showing a zero to 3-fold decrease (unpublished data). The early induction and persistence of CD4^+^ and CD8^+^ T cells may confer long-lasting memory against coronaviruses and induce durable immune responses as was observed in some SARS-CoV survivors, in whom CD4^+^ and CD8^+^ T cells persisted for 17 years (*51*). Also, IFN-γ^+^ SARS-CoV-2-specific CD8^+^ T cells have been reported in a majority of patients who recovered form COVID -19 infection (*52*).

We identified 11 SARS -CoV-2 spike-specific T cell epitopes targeted by the splenocytes of C57BL/6 mice vaccinated with SpFN+ALFQ. The most dominant and immunogenic SARS -CoV-2 spike epitope begins in the RBD domain (VNFNFNGL; aa 539 -546). This epitope was further confirmed as an MHC class I restricted K^b^(_539-546_) spike-specific CD8^+^ T cell epitope with tetramer staining. Similar to the results from splenocytes, the lungs from the SpFN+ALFQ group also showed a 3-fold increase in cells positive for the MHC class I, K^b^, restricted epitope, spike(_539-546_), compared to lungs from the SpFN+AH group. SpFN+ALFQ induced expansion of antigen specific polyfunctional CD8^+^ T cells and significantly greater amounts of polyfunctional spike -specific CD8^+^ T cells co-expressing either 3 (IFN-γ, IL-2, and TNF-α) or 2 (IFN-γ, TNF-α) cytokines compared to the SpFN+AH. To the best of our knowledge, this study is the first to report expansion of vaccine -induced SARS-CoV-2 K^b^-spike(_539-546_) -specific polyfunctional CD8^+^T cells in mice that also exhibited increased killing of peptide-pulsed target cells in an *in vitro* cytotoxic T cell assay. Since this epitope is also present in SARS -CoV, it is tempting to hypothesize that generation of cross-reactive T cells may provide protection against other coronavirus strains and warrants further investigation. The molecular mechanisms involved in specific gene signatures associated with an early induction of specific innate immune responses directly as a result of the type of vaccine formulation, which then influence the adaptive immune responses remains to be explored.

By employing a strategy of sampling immunologically relevant tissues temporally and spatially proximal to vaccination, we show that the SpFN vaccine potentiates innate sensing and mobilizes the cellular drivers of a multifactorial immune response, in particular CD4^+^ T cell responses and durable effector -memory functional CD8^+^ T cell responses, with a distinct bias towards a T_H_1 phenotype, with IFN-γ and TNFα as the dominant cytokines.

## MATERIALS AND METHODS

### Study design

The major aim of the study was to utilize a unique liposomal adjuvant formulation and a well-known and widely used adjuvant, aluminum hydroxide gel, with a novel particulate antigen, SpFN, to understand the early innate immune responses at the site of vaccination. The influence of the adjuvant formulation on the innate responses and the subsequent effects on downstream adaptive immune responses were studied in C57BL/6 mice. We utilized flow cytometry with specific markers of phenotype and activation to identify APCs and T cells at early and late time points following vaccination. Identification of SARS-CoV-2 spike-specific CD8 T cells by ELISpot analysis followed by confirmation with MHC class I -spike peptide tetramers, allowed us to evaluate the tissue distribution of antigen-specific CD8^+^ T cells for polyfunctional cytokine expression and cytolytic responses. We utilized a sample size of 5 mice/time point/vaccine formulation to obtain reliable data to perform statistical analyses and ensure reproducibility. The size was based on results of preliminary data obtained during the antigen development process. Endpoints were set based on the two major experimental arms, single immunization (endpoint Day 10) or prime/boost strategy (endpoint Day 42). All experiments were replicated in at least two to three independent experiments. The number of individual replicates for each experiment is indicated in the figure legends. Mice were age- and sex-matched and were randomly assigned to study groups. The APC flow cytometry and ELISpot analysis were performed by the investigators in a blinded fashion as they were not aware of which treatment group belonged to which adjuvant formulation. All other analyses were performed unblinded because mice were grouped according to the treatment.

### Mice vaccination and tissue processing

Female C57BL/6 mice (5-6 weeks of age) were obtained from The Jackson Laboratory. Animals were housed in groups and fed standard chow diets. Animal studies were carried out in accordance with the recommendations in the Guide for the Care and Use of Laboratory Animals of the National Institutes of Health. The protocols were approved by the Institutional Animal Care and Use Committee at the Walter Reed Army Institute of Research [Assurance number D16-00596 (A4117-01)]. Mice were vaccinated through the intramuscular route with SARS-CoV-2 spike protein (SpFN) mixed with aluminium hydroxide gel (n =5 mice/time point) or SpFN formulated with Army Liposome Formulation containing QS21 (ALFQ) (n = 5 mice/time point). Mice received either a single intramuscular vaccination in the left hind leg quadriceps muscle or the priming vaccination followed by a boost at week 3 in the right hind leg quadriceps muscle. Mice were euthanized on days 3, 5, 7, and 10 post first vaccination or 3 weeks after the second vaccination on week 6. Time points 3, 5, and 10 were repeated twice, each time with an n=5 mice/time point/adjuvant. The 6 -week time point with SpFN-ALFQ vaccine was repeated twice (n=5, n=9). Pre-vaccination peripheral blood was obtained on day -4. The inguinal and popliteal lymph nodes draining the vaccinated site, as well as, the spleen, and blood were collected at days 3, 5, 7, and 10 following the first vaccination. At the 6 -week time point, perfused lung tissue, mediastinal lymph nodes, spleen and peripheral blood were obtained.

Serum was separated from pre and post vaccination peripheral blood samples and individually stored in aliquots at -80°C until analysis. The inguinal and popliteal lymph nodes draining the vaccinated leg was combined for analysis. Lymph nodes were manually homogenized between the frosted ends of glass slides to obtain a single cell suspension. Mononuclear cells were isolated by centrifugation over Fico/Lite -LM (Atlanta Biologicals) density gradient. Single cell suspensions were then washed and resuspended in PBS with 2% FBS. Individual mouse spleens were pressed through a 70μm cell strainer using the plunger of a 3-mL syringe, to obtain a single-cell suspension, followed by washing twice in PBS containing 2 % Fetal Bovine Serum (FBS, Invitrogen). Cells were then counted using trypan blue exclusion, cryopreserved in freezing media (90% heat inactivated FBS and 10% dimethyl sulfoxide) and stored in liquid nitrogen until use.

Following mouse euthanasia, lungs were perfused with approximately 7-10 mL of 0.5 mM EDTA in Hanks Balanced Salt Solution. Lungs were cut into small pieces (approximately 3 -5 mm) and placed in a 15 mL conical tube containing 5 mL RPMI supplemented with 150 U/mL collagenase, 25 U/mL DNase IV (Fisher Scientific) and incubated for 60 minutes at 37° C on a rotator at 12-15 RPM. Tissue was then pressed through a 100μm cell strainer using the plunger of a 5-mL syringe, to obtain a single-cell suspension. Cells were washed twice in RPMI-1640 containing 10% FBS, cryopreserved in freezing media, and stored in liquid nitrogen until use.

### Protein and liposome vaccine formulation

SARS-CoV-2 prefusion antigen construct, derived from the Wuhan -Hu-1 genome sequence was designed as ferritin-fusion recombinant protein for expression as a nanoparticle. The *Helicobacter pylori* ferritin molecule was linked to the C-terminal region of pre-fusion stabilized ectodomain (residues 12 -1158). Several modifications were introduced. The spike ectodomain was modified to introduce two prolines residues (K986P, V987P) as previously described(*16*), a “GSAS” substitution at the furin cleavage site (residues 682–685), and increase the coiled-coil interactions of the spike stalk region (residues 1140-1161). The modifications were introduced to stabilize spike trimer formation on the ferritin molecule. The protein was transiently expressed as a soluble recombinant protein in mammalian Expi293 cells (Thermo Fisher Scientific). The secreted protein self -assembles into a protein nanoparticle, with the SARS -CoV-2 antigen of interest displayed on the nanoparticle surface. The culture supernatant was harvested four days post-transfection and purified by *Galanthus nivalis* lectin (GNA)-affinity chromatography and size -exclusion chromatography. Purified protein referred to as spike protein ferritin nanoparticle (SpFN) was formulated in PBS with 5% glycerol at 1 mg/mL and was subsequently diluted with dPBS to provide 10 µg per 50 µL mouse vaccination dose. The SpFN immunogen is composed of antigens from the spike protein of SARS-CoV-2 scaffolded within a ferritin nanoparticle structure. SpFN immunogen uses 24 copies of antigen spike protein domains supported in a ferritin nanoparticle vaccine platform scaffold that displays the multivalent array on its surface.

Dimyristoyl phosphatidylcholine (DMPC) and dimyristoyl phosphatidylglycerol (DMPG) saturated phospholipids, cholesterol (Chol), and synthetic monophosphoryl lipid A (MPLA, 3D -PHAD^®^) were purchased from Avanti Polar Lipids. DMPC and Chol were dissolved in chloroform, and DMPG and 3D -PHAD^®^ were dissolved in chloroform:methanol (9:1). Alhydrogel ^®^, aluminum hydroxide (AH) in a gel suspension was purchased from Brenntag. QS21 saponin was purchased from Desert King International and was dissolved in Sorensen PBS, pH 6.2 and filtered. For vaccine preparations adjuvanted with ALFQ (ALF containing QS21), lipids were mixed in a molar ratio of 9:1:12.2:0.114 (DMPC:DMPG:Chol: 3D-PHAD^®^), dried by rotary evaporation followed by overnight desiccation, rehydrated by adding Sorensen PBS, pH 6.2, followed by microfluidization to form small unilamellar vesicles (SUV) and filtration (*19, 53*). QS-21 was added to SUV to form ALFQ (DMPC:DMPG:Chol:MPLA:QS -21; 9:1:12.2:0.114:0.044).

For vaccine preparations adjuvanted with ALFQ, SpFN (600 µg/mL) was mixed with ALFQ (1.5X) in a 1:2 volume ratio. The vial was vortexed at slow speed for 1 min and then put on a roller for 15 mins. The vials were stored at 4°C for 1 hour prior to vaccination and then injected into animals within an hour. For vaccine preparations adjuvanted with AH, Alhydrogel was diluted from a stock concentration of 10 mg/mL to 900 µg/mL (1.5X) with DPBS in a sterile glass vial; SpFN (600 µg/mL) was added in a 1:2 volume ratio to the diluted Alhydrogel. The vial was vortexed at slow speed for 5 min and stored at 4°C for 1 hour prior to vaccination. Each vaccine dose in 50 μl volume contained 10 μg SpFN, 20 μg 3D-PHAD, and 10 μg QS21 (ALFQ) or 10 μg SpFN and 30 µg AL^3+^ (AH).

### Flow cytometric phenotyping and intracellular cytokine staining (ICS)

Spectral flow cytometry was performed to analyze APCs and T cells in the lymph nodes as indicated. Single cell suspensions were treated with Fc Block (BD Biosciences) for 5 minutes at 4°C before further staining. Live dead blue dye (Invitrogen) was used to stain dead cells. Analysis of APC and T cell phenotypes were performed using the fluorochrome-conjugated antibodies in table S1. Intracellular cytokine analysis was performed to identify SARS-CoV2 spike-specific T cells. Single cell suspensions from the draining lymph node were plated at 0.5×10^6^ cells per well in a 96 -well plate and stimulated with the spike (S) 1 or S2 peptide pools from JPT at a final concentration of 1 μg/mL for 6 hours at 37°C in RPMI and 10% FBS. To prevent the secretion of cytokines, GolgiStop (BD Biosciences) was added 1 hour after the addition of peptides. Spike -specific T cells were measured by surface staining followed by fixation and permeabilization using BD fixation/permeabilization solution kit (BD Biosciences,) and intracellular cytokine staining (table S1). As negative and positive controls, cells were cultured in stimulation media without peptide stimulation or with phorbol 12-myristate 13-acetate (PMA) and ionomycin (BioLegend), respectively. T cells were considered to be responsive to peptide stimulation if the frequency of cytokine-expressing T cells was greater than 0.01% of the parent population after background subtraction. Samples were analyzed on a 5 -laser Cytek Aurora flow cytometer (Cytek Biosciences). The data were analyzed using FlowJo software v10 (Tree Star, Inc.). Gating strategy for APC subsets, T cell phenotyping and ICS are shown in **fig. 1SA and fig. 2SC**, respectively.

Cryopreserved splenocytes were quickly thawed and added to 10 mL of complete RPMI 1640 media supplemented with 5% FBS and 1% Pen-strep followed by viability assessment by trypan blue exclusion method. Approximately, 1×10^6^ splenocytes were cultured in the presence of peptide pools directed towards SARS CoV -2 spike protein S1 or S2 (JPT) (1µg/mL) at 37°C, 5% CO _2_. After 1 hour of incubation with peptides, protein transport inhibitor (BD Golgi Plug™ containing Brefeldin A, 1 µg/mL and BD Golgi Stop™ containing monensin, 1 µg/mL, BD Biosciences) was added for an additional 5 hours at 37°C, 5% CO_2_. For the positive control, cells were stimulated with PMA and ionomycin (Sigma -Aldrich; 50 ng/mL and 1 μg/mL final concentration, respectively) or concavalin A (positive control for IL -4), while media served as a negative control. After the incubation period, cells were stained with LIVE/DEAD Fixable Aqua Dead Cell Stain Kit (Invitrogen), followed by surface staining with antibodies specific for the following cell surface markers (BUV737 anti -CD3, BUV395 anti-CD4, BV711 anti-CD8 (table S4), obtained from either BD Biosciences, Thermo Fisher Scientific or Biolegend, washed twice with FACS buffer and then fixed/permeabilized for 40 mins at 4°C in the dark using the eBioscience™ Intracellular Fixation & Permeabilization Buffer Set (Thermo Fisher Scientific) as per the manufacturer’s instructions. Cells were then incubated with a panel of intracellular antibodies specific for the following cytokines (V450 anti -IFN-γ, FITC anti-TNF-α, PerCP-Cy5 anti-IL-4, PE anti IL-2; table S4) for 30 min at 4°C, washed twice, and resuspended in FACS buffer. Appropriate single -color compensation controls and Fluorescence minus one (FMO) control were prepared simultaneously and were included in each analysis. Flow cytometric analysis was performed on an BD LSR II flow cytometer, and data were acquired using Diva software (BD Biosciences). The results were analyzed using FlowJo software (TreeStar). Evaluation of co-expression of different cytokines was performed using the FlowJo Boolean gate platform. The gating strategy applied for the evaluation of flow cytometry -acquired data is provided in fig. S5.

### Measurement of cytokine levels in the serum and spleen cell culture supernatants

Cytokine levels were measured using V-Plex Plus Multi -Spot Assay plates, from Meso Scale Discovery (MSD, Rockville, MD). We used the mouse Pro-inflammatory panel containing IFN -γ, IL-1β, IL-2, IL-4, IL-5, IL-6, KC/GRO (CXCL1), IL -10, IL-12p70, and TNF-α. The kit included diluent, wash buffer, detection antibody solution and read buffer, as well as calibrators and controls for each analyte, from the manufacturer. Plates were washed three times with MSD wash buffer before the addition of MSD reference standard and calibrator controls used for quantifying antibody concentrations. Serum samples were diluted 1:2 in MSD diluent buffer, then added to the wells in duplicate. Plates were incubated overnight at 4°C, then washed three times. MSD detection antibody solution was added to each well, plates were incubated for 2 hours at RT with shaking at 500 rpm then washed three times. MSD 2x read buffer T was added to each well. Plates were read by MESO SECTOR S 120 Reader. Analyte concentration was calculated using DISCOVERY WORKBENCH^®^ MSD Software and reported as picograms/mL. For cell supernatants, approximately 1×10^6^ cells were cultured in the presence of peptide pools directed towards SARS CoV-2 spike protein (JPT, 1µg/mL) for 6 hours at 37°C, 5% CO_2_. Cells were centrifuged at 500 g for 10 minutes and the supernatant was collected. Supernatants were diluted 1:2 in MSD diluent buffer and subsequently processed as described above.

### ELISpot

Multiscreen plates (Millipore, Bedford MA) were coated with anti-IFN-γ antibody (capture mAb; 1 µg/mL) according to the manufacturer’s instructions (R&D Systems, Minneapolis, MN). Plates were blocked using culture medium (DMEM containing 10% FBS, Pen/Strep, HEPES, NEAA, sodium pyruvate, 2-mercaptoethanol). Thawed splenocytes were counted and plated at 4 x 10^6^ cells/mL (50 μL/well) in triplicate. Recombinant peptides, Epitope Mapping Peptide Set (EMPS), SARS-CoV-2, Spike protein (JPT Peptide Technologies, Berlin, Germany), tested at 1.25 µg/mL and anti-CD3 (as positive control) were used to stimulate the cultures. Plates were incubated for 42 hrs in the presence of antigen at 37°C and then cells were removed by washing the plates in an automated plate washer. Plates were incubated with biotinylated anti -IFN-γ antibody (detecting mAb; 1 µg/mL) overnight at 4°C. Plates were washed and incubated with Streptavidin -alkaline phosphatase for 2 hrs at RT. Color development was done using ELISpot Blue Color Module (R&D Systems, Minneapolis, MN) and processed according to the manufacturer’s instructions. Plates were counted and data analyzed using the AID Autoimmun Diagnostica GmbH ELISpot reader (Strassberg, Germany) and software. T cell responses were considered positive when the mean spot count exceeded the mean ± 3 SD of the negative control wells.

### Enumeration of SARS CoV-2 Spike-specific CD8^+^ T cells by tetramer staining

MHC Class I or H-2K^b^ restricted SARS-CoV-2 spike peptide (aa 539-546; VNFNFNGL)-PE conjugated tetramer (NIH tetramer core facility) was used for the detection of antigen specific CD8^+^ T cells. (see table S2 for a list of tetramers used in the assay). Approximately, 1×10^6^ freshly isolated splenocytes from mice were first stained with a dead cell discrimination dye (Aqua stain, 1:1000 dilution, Invitrogen) for 30 minutes at 4°C followed by washing twice with 1XPBS. Cells were blocked with anti -mouse CD16/CD32 for 30 mins followed by further incubation with 1µL (1.2µg) of PE-conjugated H-2K^b^ SARS CoV-2 spike-specific tetramer for 30 min at 4 °C in the dark. After washing, cells were further surface stained with BUV737 -anti-mouse CD3, BUV395-anti-mouse CD4, BV711-anti-mouse CD8a and incubated for an additional 30 mins in the dark at room temperature, washed twice, and resuspended in FACS buffer. Flow cytometric analysis was performed on a BD LSR II flow cytometer and data were acquired using Diva software (BD Biosciences). The results were analyzed using FlowJo software (TreeStar). Appropriate single-color compensation controls and fluorescence minus one (FMO) control were prepared simultaneously and were included. Staining with H -2K^d^ restricted SARS-CoV-2 spike peptide (aa 365-373; CYGVSPTKL)-PE conjugated tetramer served as a control tetramer in this analysis.

### Detection of SARS-CoV-2 spike-specific (Tetramer positive) CD8^+^ T cells secreting cytokines by flow cytometry

To detect the epitope specific CD8^+^ T cell cytokine production, we combined antigen stimulation with tetramer labelling and intracellular cytokine staining. Splenocytes (2 × 10^6^/well) were cultured in a 96 -well v-bottom plate in the presence of peptide pools directed towards SARS CoV -2 spike protein S1 (1µg/mL) at 37°C, 5% CO_2_. After 1 hour of incubation with peptides, protein transport inhibitor (BD Golgi Plug™ containing Brefeldin A, 1 µg/mL and BD Golgi Stop™ containing monensin, 1 µg/mL, BD Biosciences) was added followed by an additional 5-hour incubation at 37°C, 5% CO_2_. At the end of the incubation period, cells were washed at 300 g at 4°C for 7 mins, then labeled with a dead cell discrimination dye (Aqua stain, 1:1000 dilution, Invitrogen) for 30 mins at 4°C followed by washing twice with 1XPBS. Cells were fixed/permeabilized for 40 mins at 4°C in the dark using the eBioscience™ Intracellular Fixation & Permeabilization Buffer Set as per the manufacturer’s instructions. Cells were then blocked for 20 min at 4°C in Fc -blocking solution (5 μg/mL of CD16/CD32 mAb, eBioscience, USA) and further stained for 40 mins at RT with 1µL (1.2µg) MHC class I or H -2K^b^ restricted SARS-CoV-2-spike peptide (aa 539-546; VNFNFNGL)-PE conjugated tetramer (NIH tetramer core facility) diluted in Perm/wash buffer. In the last 20 min of tetramer incubation, the following mix of fluorescent antibodies were added: BUV737-anti-mouse CD3, BUV395-anti-mouse CD4, BV711-anti-mouse CD8a and antibodies directed towards intracellular cytokines such as V450 anti -IFN-γ, FITC anti-TNF-α, and APC anti IL-2 (**Supplementary Table 4**), then washed twice and resuspended in FACS buffer. Appropriate single -color compensation controls and Fluorescence minus one (FMO) control were prepared simultaneously and were included in each analysis. Flow cytometric analysis was performed on a BD LSR II flow cytometer and data were acquired using Diva software (BD Biosciences). The results were analyzed using FlowJo software (Tree Star). Staining with H-2K^d^ restricted SARS-CoV-2 Spike peptide (aa 365-373; CYGVSPTKL)-PE conjugated tetramer served as a control tetramer in this analysis.

### Cytotoxic T lymphocyte killing assay

Single cell suspensions from spleens of vaccinated mice were used as effector cells and naive spleens were used as target cells. Target cells were prepared by diluting cells to 20×10^6^/mL and making 2 x 500 µL aliquots (10×10^6^ each). One aliquot was incubated in the presence of SARS CoV -2 spike protein specific peptides LVKNKCVNFNFNGLT and KCVNFNFNGLTGTGV at 1 µg/mL (JPT) and the other aliquot was incubated in media for 45 min in a 37°C water bath. Both aliquots were washed twice with 3 mL PBS. CFSE high and low solutions were made at 0.5 and 0.05 µM respectively using ThermoFisher CellTrace™ CFSE Cell Proliferation Kit (Catalog C34554). Peptide pulsed and non-pulsed cells were resuspended respectively in CFSE high and low solutions, incubated for 10 min in a 37°C water bath, and then washed twice in media. Target cells were resuspended at 2×10^6^/mL and 50 µL of each CFSE high and low (1:1, CFSE high:CFSE low) was added to each well. Effector cells were resuspended at 40×10^6^/mL (10×10^6^ cells in 250 µL) and 2 -fold serial dilutions were made by adding 125 µL of cells to 125 µL of media. 100 µL of each dilution was added to a well containing the target cell mixture, bringing the total volume to 200 µL. Cells were co -incubated overnight at 37°C, 5% CO_2_. Following incubation, cells were stained with Live/Dead Aqua stain (1:1000 dilution) (Invitrogen, Thermo Fisher Scientific) for 30 minutes at 4°C followed by washing twice with 1XPBS. Then, the cells were blocked with anti -mouse CD16/CD32 for 30 mins followed by further incubation with 1µL (1. 2µg) of PE-conjugated H-2K^b^ spike_(539-546)_-specific tetramer for 30 min at 4 °C in the dark. After washing, cells were further stained with BUV737-anti-mouse CD3, BUV395-anti-mouse CD4, BV711-anti-mouse CD8a and incubated for an additional 30 mins in the dark at room temperature. Flow cytometric acquisition and data analysis were performed as described above with appropriate controls.

### Statistical analysis

Flow cytometric data were analyzed using FlowJo v.10.0.8 (BD). Data are displayed as dot plots, or bar graphs. AH and ALFQ groups were compared by unpaired Student’s t-test or Mann–Whitney U-test or Pearson correlation. Graphs were plotted using GraphPad Prism v.8.4.0. Statistical analyses were conducted using GraphPad Prism v.8.4.0 software. P values ≤0.05 were considered statistically significant.

## Acknowledgments

Mouse anti-RBD monoclonal antibody was produced under HHSN272201400008C and obtained through BEI Resources, NIAID, NIH: Spike Glycoprotein Receptor Binding Domain (RBD) from SARS -Related Coronavirus 2, Wuhan-Hu-1 with C-Terminal Histidine Tag, Recombinant from HEK293F Cells, NR -52366. SARS CoV-2 MHC class I tetramers were obtained from NIH tetramer core facility at Emory, Atlanta.

## Author Contributions

Conceptualization and study design: J.M.C., S.S., J.R.C., E.B -L., A.M.W.M., and M.R. Methodology: J.M.C., S.S., Z.L., E.B-L., A.A., G.R.M., R.S.S., W-E.C., W.C.C., M.G.J., K.M., E.B.M., and. J.S.B. Data analysis: J.M.C., S.S., Z.L., E.L-B., J.R.C., A.M.W.M., and M.R. Visualization: J.M.C., S.S., Z.L., E.B-L. Supervision: M.R. and A.M.W.M in their respective institutions. Funding acquisition: K.M and N.L.M. Intellectual input: N.L.M. Project administration: K.M. and N.L.M. Writing (original draft): J.M.C., S.S., J.R.C., E.B-L., Z.L., A.M.W.M., and M.R. Writing (review and editing): All authors.

## Data availability

All data needed to evaluate the conclusions in the paper are present in the paper or in the Supplementary Figs. 1 to 6. The spike (S) protein sequence was derived from the Wuhan-Hu-1 genome sequence (GenBank accession number: MN908947.3).

## SUPPLEMENTARY MATERIALS

**Fig. S1.**
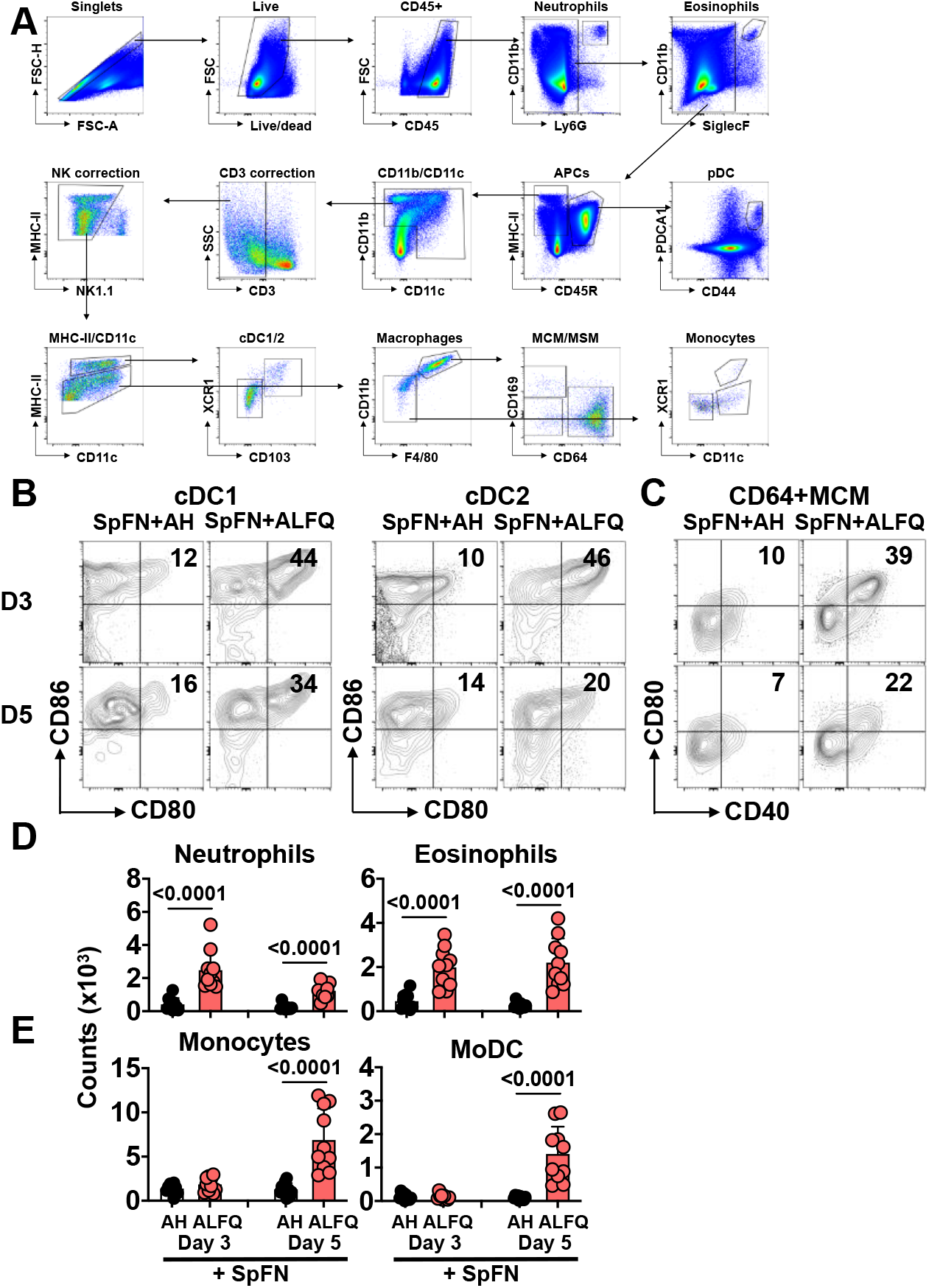
Flow cytometry analysis of innate immune cells in draining lymph nodes responding to SpFN vaccine adjuvanted with ALFQ or AH. **(A)** Flow cytometry gating strategy for the identification of neutrophils (Ly6G^+^CD11b^+^), eosinophils (Siglec F^+^CD11b^+^), pDCs (MHC-II^+^CD45R^+^PDCA1^+^), cDC1 (MHC-II high CD11c^+^XCR1^+^CD103^+^), cDC2 (MHC-II high CD11c^+^XCR1^-^), MSM (MHC II^+^F4/80^+^CD11b^+^CD169^+^CD64^-^), MCM (MHC-II^+^F4/80^+^CD11b^+^CD169-CD64+/-), XCR1+cDC1-like moDCs (MHC-II+CD11b int CD11c^+^XCR1^+^), XCR1-CD11c^+^moDCs (MHC-II^+^CD11b int CD11c^+^XCR1^-^), and monocytes (MHC-II^+^CD11b int CD11c^-^XCR1^-^). **(B)** Flow cytometry plots demonstrating expression of costimulatory molecules CD80 and CD86 by cDC upon activation after vaccination. (**C)** Flow cytometry plots demonstrating activation of macrophages by expression of CD40 and CD80. **(D)** Magnitude of non-classical APCs, including neutrophils, eosinophils, **(E)** XCR1^+^moDCs, and monocytes at days 3 and 5 post priming vaccination.

**Fig. S2.**
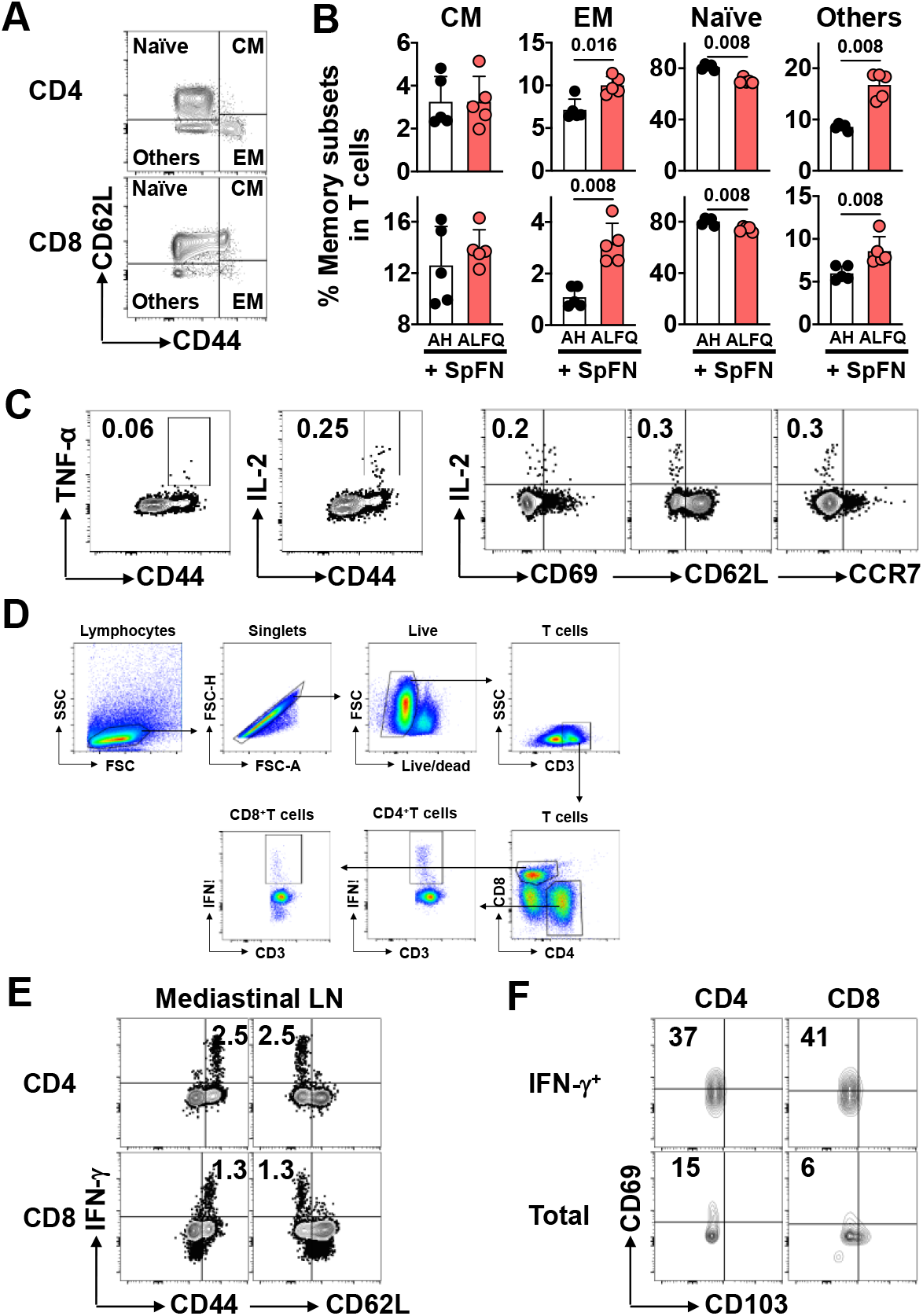
Characterization of the responding T cells upon priming with SpFN vaccine. **(A)** Memory differentiation phenotypes shown as flow plots (left panels) and quantitation of phenotype differences between the two vaccine groups at day 7 after the first vaccination (right panel). **(B)** Memory T cells are determined as naive T cells (CD62L^+^CD44^-^), central memory T cells (CD62L^+^CD44^+^), effector memory T cells (CD62L^-^ CD44^+^), and others. **(C)** Cytokine expression and activation markers characterizing SARS -CoV-2 spike-specific CD4^+^ T cells at day 10 day after the first vaccination. **(D)** Flow cytometry gating strategy for the identification of SARS-CoV-2 spike-specific CD4^+^ and CD8^+^ T cells in the mediastinal lymph nodes. (**E-F)**, Characterizing the SARS-CoV-2 spike-specific CD4^+^ and CD8^+^ T cells in the mediastinal lymph nodes at 3 weeks following prime-boost vaccination by the expression of IFNα. IFNα^+^T cells are **(E)**, CD44^+^CD62L^-^ and **(F)** more frequently CD69^+^ than parent populations but CD103^-^, indicating effector memory and activation. Bars indicate mean ± s.d. Differences between the two groups were analyzed by using non -parametric Mann–Whitney U-test with P ≤ 0.05 considered as statistically significant.

**Fig. S3.**
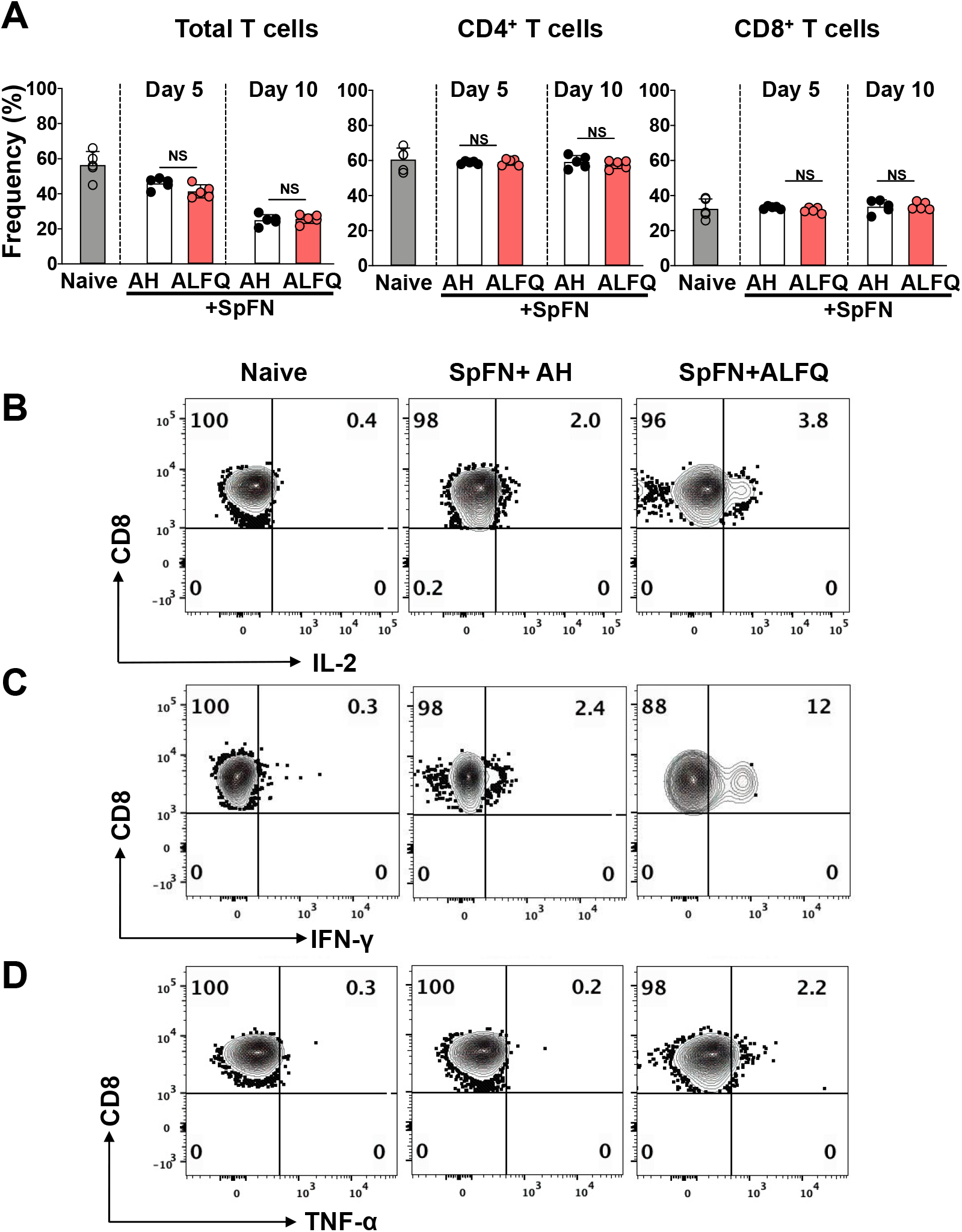
Frequency of T cells in splenocytes of mice following priming vaccination with SpFN vaccine. **(A)** Frequency of total CD3^+^, CD4^+^ and CD8^+^ T cells in splenocytes of mice on days 5 and 10 post priming vaccination with SpFN adjuvanted with AH or ALFQ. **(B-D)** Representative flow plots depicting the frequency of (**B)** IL-2 **(C)** IFN-γ and (**D)** TNF-α secreting CD8^+^ T cells on day 10 post priming vaccination.

**Fig. S4.**
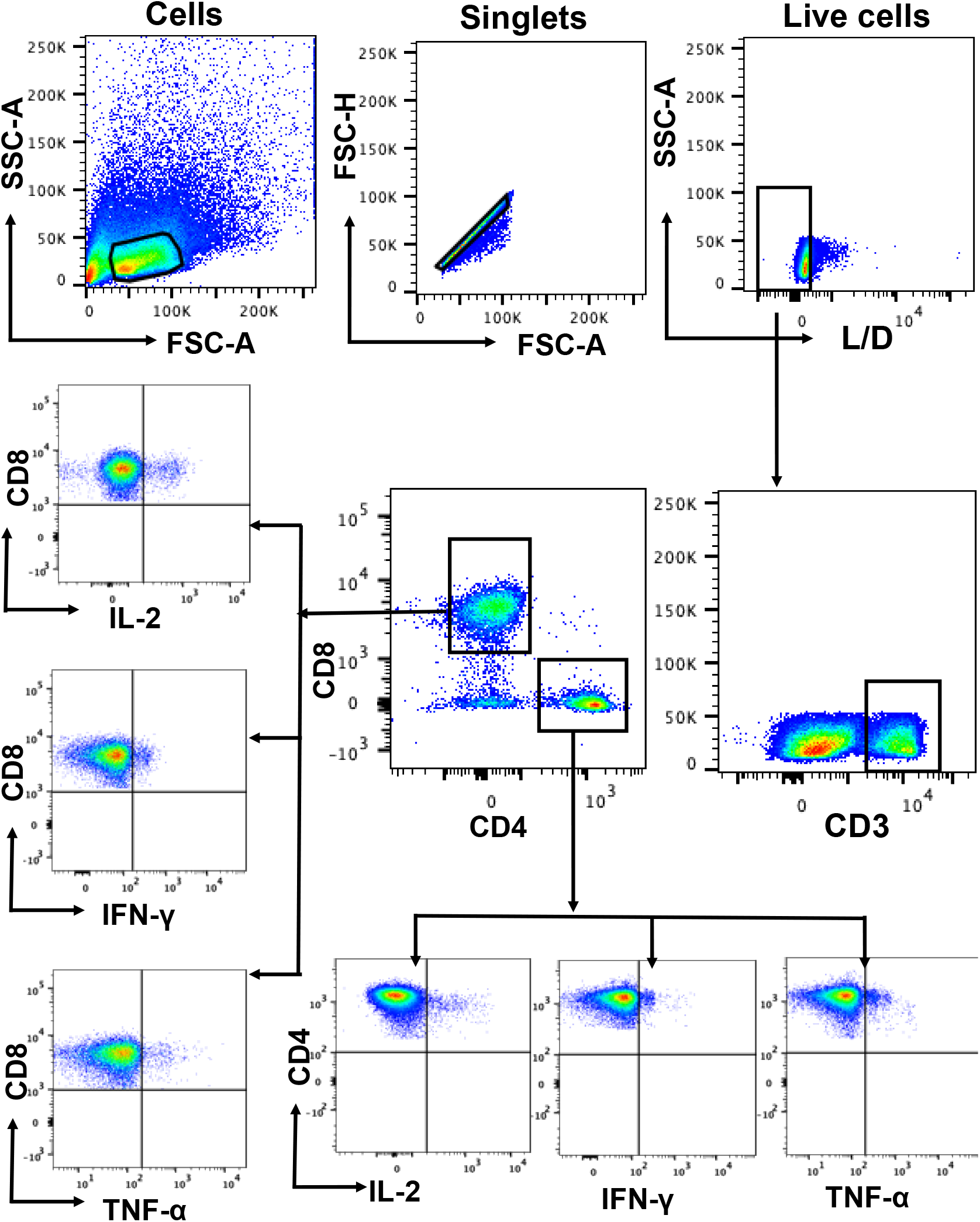
Flow gating strategy to determine the frequency of cytokine secreting CD4^+^ and CD8^+^ T cells. Flow gating strategy to determine the frequency of cytokine secreting IL -2, IFN-γ, and TNF-α secreting CD4^+^ and CD8^+^ T cells. Gates were defined on the respective fluorescence minus one (FMO) control.

**Fig. S5.**
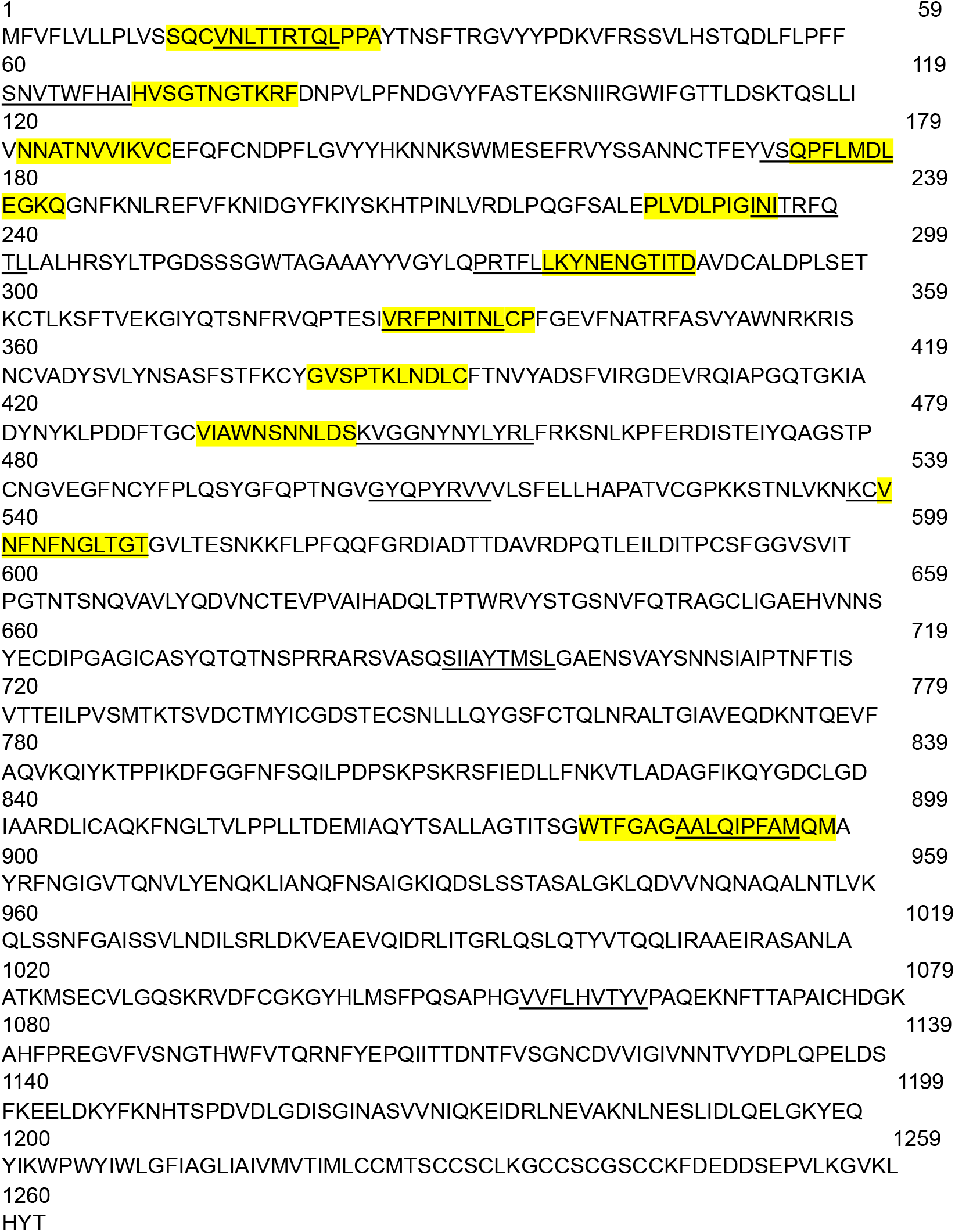
Amino acid sequence of the SARS-CoV-2 spike protein. Highlighted peptides were part of ELISpot -reactive SARS-CoV-2 glycoprotein derived matrix peptide pools. Underlined peptides are sequences predicted by NETMHC prediction algorithm (iedb.org). Numbers indicate amino acid residues.

**Fig. S6.**
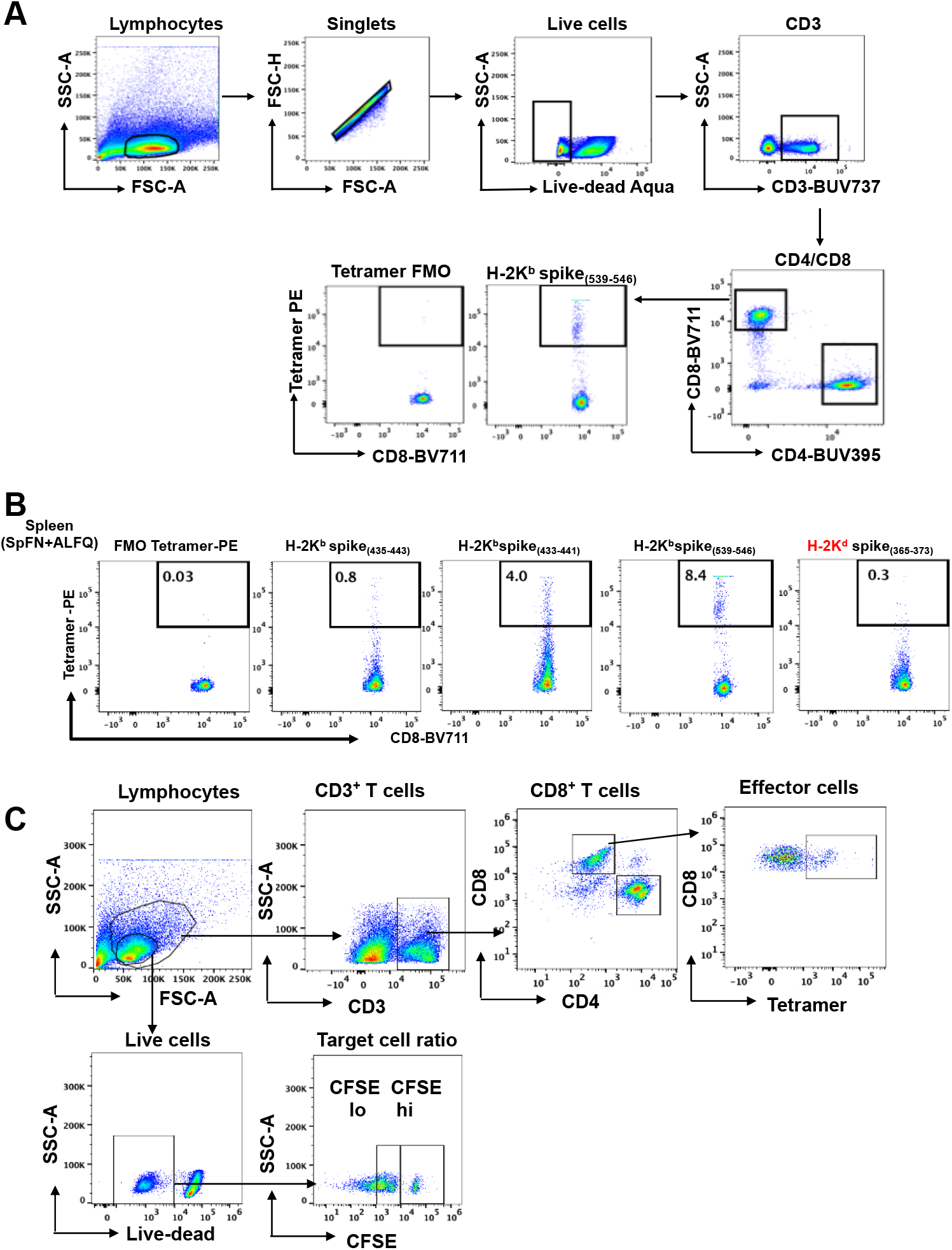
Flow gating strategy for determining the frequency of SARS -CoV-2 specific CD8^+^ T cells and for the CTL assay following SpFN vaccination. **(A)** Flow gating strategy to determine the frequency of SARS -CoV-2 spike specific CD8^+^ T cells in the splenocytes of mice vaccinated with SpFN+AH vs. SpFN+ALFQ as detected by MHC class I tetramer staining. **(B)** Cells were stained for four different tetramers directed towards four different epitopes on SARS -CoV-2 spike protein [K^b^ _(433-441)_; _(435-443)_; _(539-546)_ and K^d^ _(365-373)_]. Detailed information is provided in **table S3. (C)** Gating strategy used to determine the frequency of tetramer positive CD8^+^ T cells (effector cells) and ratio of CFSE high (hi): low (lo) target cells.

**Table S1.** List of antibodies used for Cytex flow cytometry based analysis of the APC phenotypes.

| Antigen | Fluorochrome | Clone | Manufacturer | Catalog number |
| --- | --- | --- | --- | --- |
| CD45R/B220 | BUV395 | RA3-6B2 | BD | 563793 |
| CD80 | BUV661 | 16-10A1 | BD | 741515 |
| CD3 | BUV737 | 145-2C11 | BD | 612771 |
| CD86 | BUV805 | GL1 | BD | 741946 |
| CCR7 | BV421 | 4B12 | BioLegend | 120120 |
| CD40 | BV480 | 3/23 | BD | 746970 |
| XCR1 | BV510 | ZET | BioLegend | 148218 |
| CD11c | BV570 | N418 | BioLegend | 117331 |
| CD169 | BV605 | 3D6.112 | BioLegend | 142413 |
| F4/80 | BV650 | BM8 | BioLegend | 123149 |
| I A/E | BV711 | M5/114.15.2 | BioLegend | 107643 |
| CD117 | BV750 | 2B8 | BD | 747412 |
| CD44 | BV785 | IM7 | BioLegend | 103059 |
| CXCR5 | FITC | L138D7 | BioLegend | 145520 |
| CD8a | PerCP | 53-6.7 | BioLegend | 100732 |
| CD24 | PerCP-Cy5.5 | M1/69 | BioLegend | 101824 |
| PDCA1 | PE | REA818 | Miltenyi Biotec | 130-112-220 |
| CD103 | PE-Dazzle594 | 2E7 | BioLegend | 121430 |
| CD11b | PE-Cy5 | M1/70 | BioLegend | 101210 |
| Siglec-F | PE-Vio770 | REA798 | Miltenyi Biotec | 130-112-176 |
| CD64 | AF647 | X54-5/7.1 | BioLegend | 139322 |
| Ly6G | AF700 | 1A8 | BioLegend | 127610 |
| NK1.1 | APC-Cy7 | PK136 | BioLegend | 108724 |
| CD45 | APC-Fire810 | 30-F11 | BioLegend | 103174 |

**Table S2.** List of antibodies used for Cytex flow cytometry based analysis of the T cell phenotypes.

| Antigen | Fluorochrome | Clone | Manufacturer | Catalog number |
| --- | --- | --- | --- | --- |
| CD3 | BUV395 | 145-2C11 | BD | 563565 |
| CD69 | BV421 | H1.2F3 | BioLegend | 104528 |
| CD44 | BV510 | IM7 | BioLegend | 103044 |
| CD62L | BV570 | MEL-14 | BioLegend | 104433 |
| PD-1 | BV605 | 29F.1A2 | BioLegend | 135219 |
| ICOS | BV650 | 398.4A | BioLegend | 313550 |
| CD8 | PerCP-Cy5.5 | 53-6.7 | BioLegend | 100734 |
| CXCR5 | PE | L138D7 | BioLegend | 145504 |
| CD4 | PE-Dazzle 594 | GK1.5 | BioLegend | 100456 |
| CCR7 | PE-Cy5 | 4B12 | BioLegend | 120114 |
| CD45 | APC-Fire810 | 30-F11 | BioLegend | 103174 |

**Table S3.**
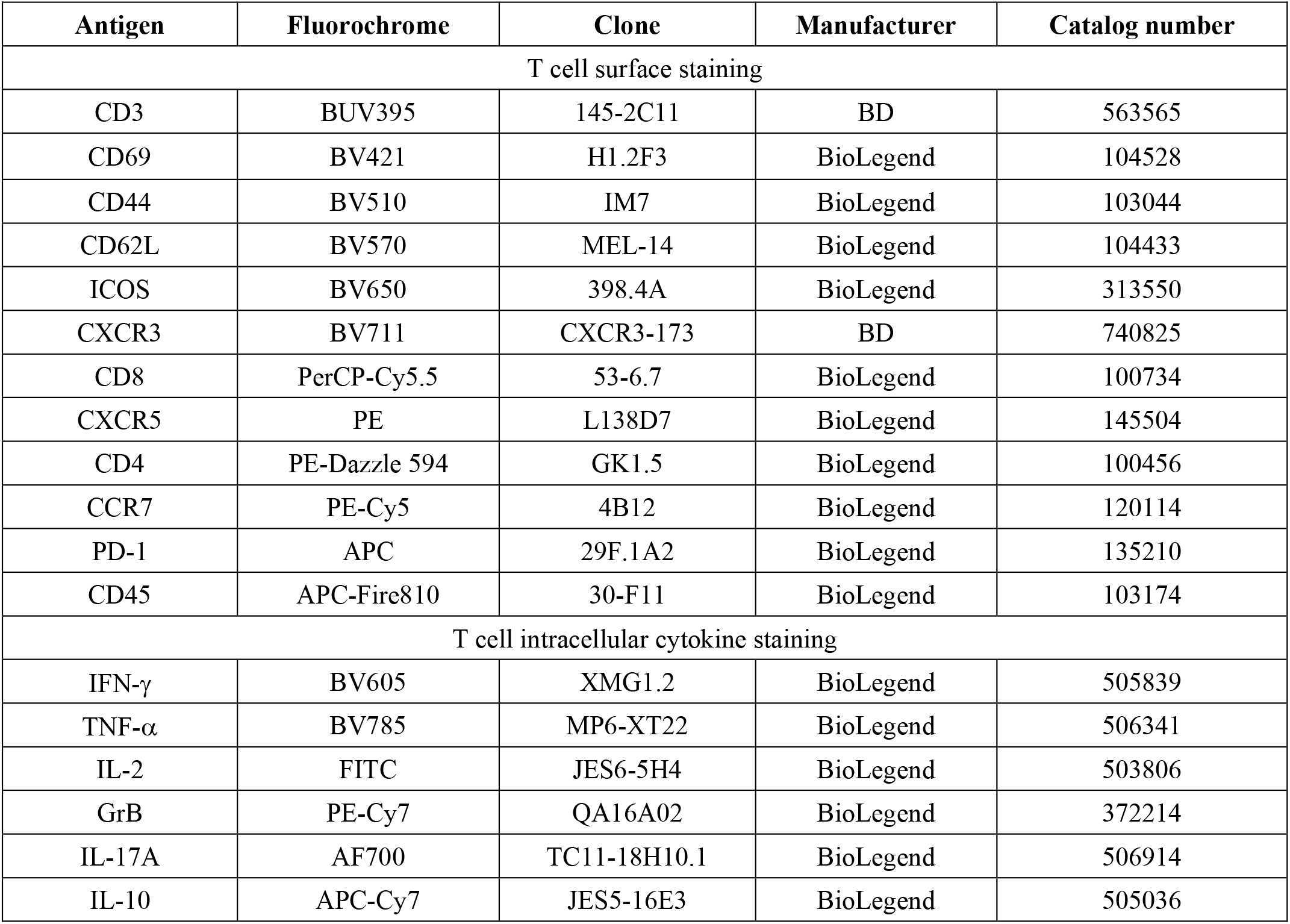
List of antibodies used for Cytex flow cytometry based analysis of the T cell cytokine staining.

| Antigen | Fluorochrome | Clone | Manufacturer | Catalog number |
| --- | --- | --- | --- | --- |
| T cell surface staining |  |  |  |  |
| CD3 | BUV395 | 145-2C11 | BD | 563565 |
| CD69 | BV421 | H1.2F3 | BioLegend | 104528 |
| CD44 | BV510 | IM7 | BioLegend | 103044 |
| CD62L | BV570 | MEL-14 | BioLegend | 104433 |
| ICOS | BV650 | 398.4A | BioLegend | 313550 |
| CXCR3 | BV711 | CXCR3-173 | BD | 740825 |
| CD8 | PerCP-Cy5.5 | 53-6.7 | BioLegend | 100734 |
| CXCR5 | PE | L138D7 | BioLegend | 145504 |
| CD4 | PE-Dazzle 594 | GK1.5 | BioLegend | 100456 |
| CCR7 | PE-Cy5 | 4B12 | BioLegend | 120114 |
| PD-1 | APC | 29F.1A2 | BioLegend | 135210 |
| CD45 | APC-Fire810 | 30-F11 | BioLegend | 103174 |
| T cell intracellular cytokine staining |  |  |  |  |
| IFN-γ | BV605 | XMG1.2 | BioLegend | 505839 |
| TNF-α | BV785 | MP6-XT22 | BioLegend | 506341 |
| IL-2 | FITC | JES6-5H4 | BioLegend | 503806 |
| GrB | PE-Cy7 | QA16A02 | BioLegend | 372214 |
| IL-17A | AF700 | TC11-18H10.1 | BioLegend | 506914 |
| IL-10 | APC-Cy7 | JES5-16E3 | BioLegend | 505036 |

**Table S4.** List of antibodies used for intracellular cytokine staining by BD LSRII flow cytometry based analysis of the cytokine response in the splenocytes.

| Antigen | Fluorochrome | Clone | Manufacturer | Catalog number |
| --- | --- | --- | --- | --- |
| CD3 | BUV737 | 145-2C11 | BD | 612771 |
| CD4 | BUV395 | GK1.5 | BD | 565974 |
| CD8 | BV711 | 53-6.7 | BD | 563046 |
| IFN-γ | V450 | XMG1.2 | BD | 560661 |
| IL-4 | PerCP-Cy5 | 11B11 | BD | 560700 |
| TNF-α | FITC | MP6-XT22 | BD | 554418 |
| IL-2 | APC | JES6-5H4 | Biolegend | 503810 |
| IL-2 | PE | JES6-5H4 | Biolegend | 503808 |

**Table S5.**
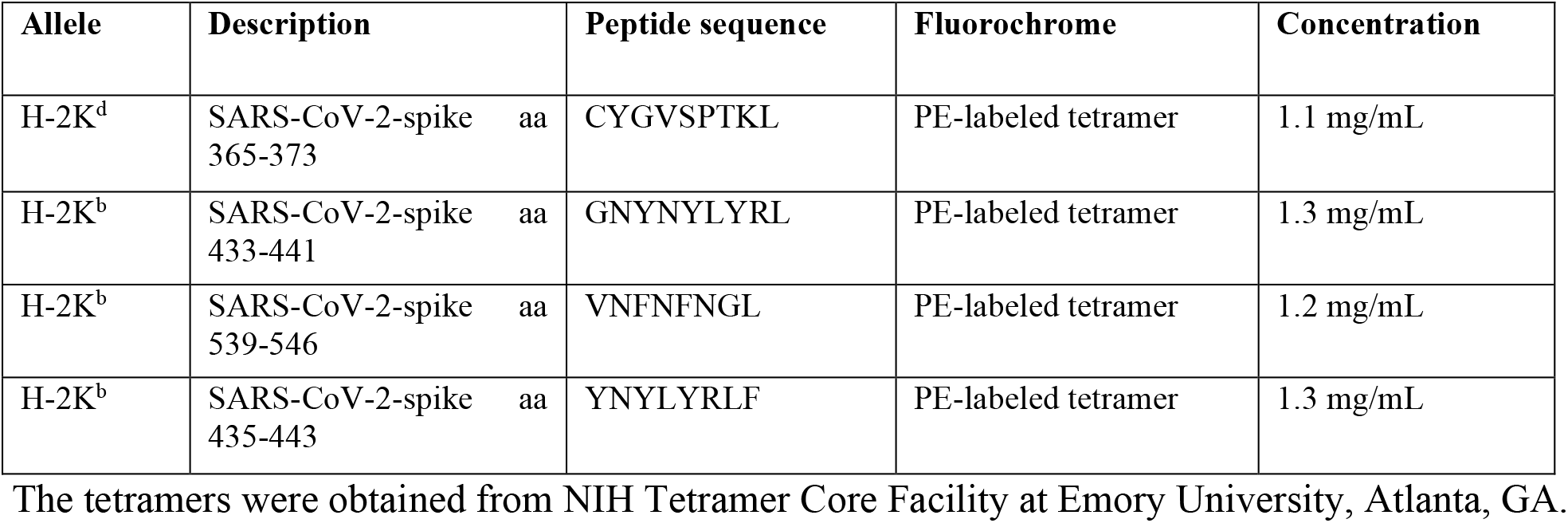
List of MHC class I tetramer used for the detection of SARS-CoV-2 spike specific CD8^+^ T cell response.

| Allele | Description | Peptide sequence | Fluorochrome | Concentration |
| --- | --- | --- | --- | --- |
| H-2K<sup>d</sup> | SARS-CoV-2-spike aa<br>365-373 | CYGVSPTKL | PE-labeled tetramer | 1.1 mg/mL |
| H-2K<sup>b</sup> | SARS-CoV-2-spike aa<br>433-441 | GNYNYLYRL | PE-labeled tetramer | 1.3 mg/mL |
| H-2K<sup>b</sup> | SARS-CoV-2-spike aa<br>539-546 | VNFNFNGL | PE-labeled tetramer | 1.2 mg/mL |
| H-2K<sup>b</sup> | SARS-CoV-2-spike aa<br>435-443 | YNYLYRLF | PE-labeled tetramer | 1.3 mg/mL |

The tetramers were obtained from NIH Tetramer Core Facility at Emory University, Atlanta, GA.

